# Omix: A Multi-Omics Integration Pipeline

**DOI:** 10.1101/2023.08.30.555486

**Authors:** Eléonore Schneegans, Nurun Fancy, Michael Thomas, Nanet Willumsen, Paul M Matthews, Johanna Jackson

**Affiliations:** UK Dementia Research Institute at Imperial College London, London W12 0BZ, UK; Department of Brain Sciences, Imperial College London, London W12 0BZ, UK

## Abstract

**Summary:** The *Omix* pipeline offers an integration and analysis framework for multiomics intended to preprocess, analyse, and visualise multimodal data flexibly to address various research questions. From biomarker discovery and patient stratification to the investigation of complex biological processes, *Omix* empowers researchers to derive valuable insights from omics data. Using Alzheimer’s Disease (AD) bulk proteomics and transcriptomics datasets generated from two distinct regions derived from post-mortem brains, we demonstrate the utility of *Omix* in generating an integrated pseudo-temporal multi-omics profile of AD.

**Availability and Implementation:** *Omix* is implemented as a software package in R. The code for the *Omix* package is available at https://github.com/eleonoreschneeg/Omix. Reference documentation and online tutorials are available at https://eleonore-schneeg.github.io/Omix. All code is open-source and available under the GNU General Public License v3.0 (GPL-3).

## 1 Introduction

Recent advancements in high-throughput omics techniques have transformed biomedical research by enabling the comprehensive capture of biological information across multiple molecular layers. This progress has led to groundbreaking discoveries in understanding biological systems. However, analysing individual omic layers offers only a partial glimpse into the complexity of these systems. Multi-omics integration unveils the interplay between molecular layers, providing comprehensive mechanistic insights into health and disease (Subramanian et al., 2020). For instance, integrating proteomic and bulk transcriptomic data provides a powerful tool for studying biological processes, identifying disease markers, and understanding dysregulated pathways. By combining information on transcription factor targets, protein abundance, and gene expression levels, researchers can gain insights into regulatory mechanisms and the functional consequences of altered gene expression in disease states.

Integrating multiple omics layers poses a significant challenge across various stages, including data processing, modelling, and the biological interpretation of results. The complexity inherent in omics data necessitates the use of diverse processing methods, often requiring the combination of multiple software tools and extensive computational expertise. More so, the rapid evolution of integration tools introduces obstacles to reproducibility and hampers the achievement of FAIR principles (Findability, Accessibility, Interoperability, and Reusability) of multi-omics efforts (Krassowski et al., 2020). Currently, this is driving the demand for more end-to-end pipelines, which will ultimately widen the adoption of multi-omics approaches within the research community.

To tackle these challenges, we developed *Omix*, an efficient R software package that processes, integrates, and analyses multi-omic data (transcriptomic and proteomic in this instance) in an end-to-end manner. *Omix* provides a wide range of cutting-edge processing functions, integrative models, and quality control features. It allows researchers to easily explore different integration strategies, enhancing the speed, scalability, and flexibility of multi-omics analyses. What sets *Omix* apart from existing R packages like Miodin (Ulfenborg, 2019) or Movics (Lu et al., 2020) is its comprehensive range of model choices and inclusion of both preand post-integration steps, along with visualisations to aid in the biological interpretation of results (Table 1). *Omix* is suitable for various research objectives, including biomarker discovery, patient stratification, and elucidating biological mechanisms, whereas other tools often focus on specific areas. The modular framework of *Omix* enables the storage of analysis parameters and results from different algorithms (both single-omic and integrative) within the same object, facilitating easy comparison of outputs. This design also allows for the incorporation of additional integrative models as the field progresses. While the current version of *Omix* primarily focuses on bulk transcriptomics and proteomics, future versions will encompass a broader range of omics types, expanding the software’s applicability and usefulness.

**Table 1.** Multi-omics integration softwares landscape. Biomarker Discovery (BD), Biological Mechanisms (BM), Sample stratification (STR)

|  | Supported omics | Pre-processing | Single omic analysis | Integrative models | Use case | Downstream analyses | Interactive visualisations | Language | Ref |
| --- | --- | --- | --- | --- | --- | --- | --- | --- | --- |
| **Miodin** | SNP, RNA, methylation, Proteins, | + | - | MOFA | BM | - | - | R | (Ulfenborg, 2019) |
| **MiBiOmics** | miRNA, RNA, Proteins | - | - | Co-inertia analysis | BD | Multi-omics networks | + | Web | (Zoppi et al., 2021) |
| **Muon** | Single cell omics | - | - | MOFA, WNN | NA | - | - | Python | (Bredikhin et al., 2022) |
| **Movics** | SNP, RNA, methylation, Proteins | - | - | Range of clustering algorithm | STR | Survival analysis, enrichment | - | R | (Lu et al., 2020) |
| **Omix** | RNA, Proteins | + | + | MOFA,MEIFESTO, sMBPLS, MBPLS, DIABLO, iCluster | BD, BM, STR | Multi-omics signatures, networks, modules, functional/ cell type/ TF enrichment, pseudotime analysis | + | R |  |

## 2 Implementation

### 2.1 Package Overview

The *Omix* pipeline offers an integration and analysis framework for multi-omics intended to pre-process, analyse, and visualise multimodal data flexibly to address research questions. *Omix* is built on four consecutive blocks, (1) preparation of the multimodal container, (2) processing and quality control, (3) single omic analyses, and (4) multi-omics vertical integration, presented in Figure 1.

**Figure 1.**
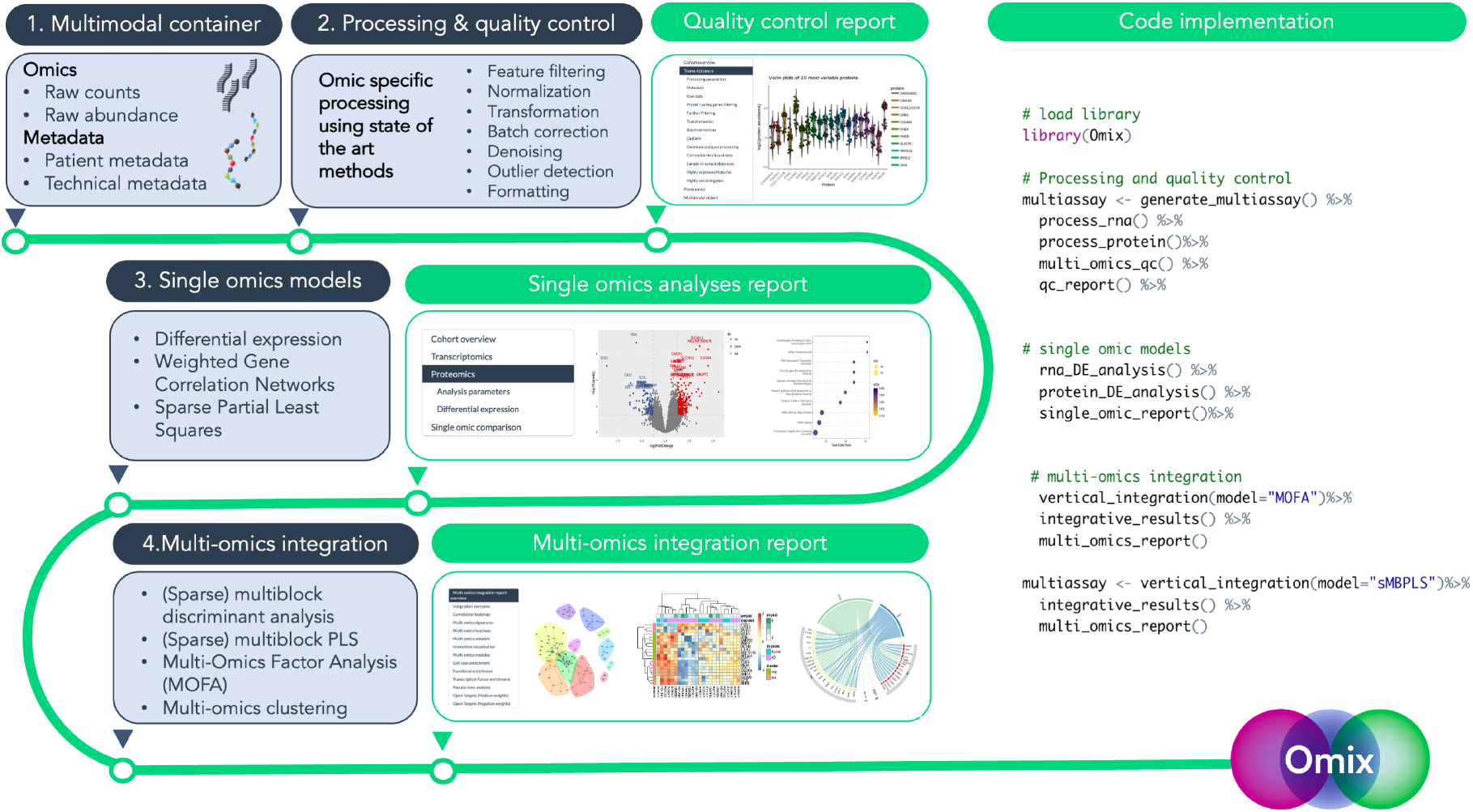
The Omix pipeline. The pipeline is organised around four building blocks, producing standardised outputs that can be visualised in interac:ve reports.

The *Omix* R package is stored in a publicly accessible Github repository with version control. To ensure reproducibility, continuous integration (CI) Github Actions perform build and functionality tests, as well as rebuild package vignettes using a small test dataset after updates. Upon successful completion, a version-tagged Docker image is generated using a Docker file that installs required software and system-level dependencies. This enables users to conduct end-to-end reproducible analyses without explicitly configuring package dependencies. The Docker image is uploaded to a Docker registry named *Omix*.

### 2.2 Multimodal container

The *Omix* multimodal container uses the Bioconductor *MultiAssayExperiment* format, a widely recognised format for managing multiple omics datasets and associated metadata (Ramos et al., 2017). The MultiassayExperiment serves as a comprehensive storage unit for all raw and processed data, as well as the associated parameters for processing and analysis, quality control metrics, and results.

### 2.3 Processing and quality control

Each omics layer is independently processed using established protocols, such as DESeq2 for bulk transcriptomics (Love et al., 2014), or Probatch for proteomics (Čuklina et al., 2021). Users have the flexibility to choose which parameters and steps of the modular sequence they want to execute, based on their analysis requirements. The available options include feature filtering, normalisation, transformation, batch correction, denoising, and sample outlier removal. Furthermore, automatic ontology detection enables the translation of features between different ontologies, such as a single gene name from Ensembl ID or UniProt ID, thereby facilitating the interpretation of results.

### 2.4 Single omic models

*Omix* presents a range of analytical tools, such as Differential Analysis (DE), which is a conventional method for identifying transcripts/proteins that display differential expression between distinct grouping variables or with regression. Additionally, the Weighted Gene Correlation Network Analysis (WGCNA) is available to identify gene modules that are associated with specific disease covariates (Langfelder & Horvath, 2008). The Sparse Partial Least Squares (sPLS) method is also offered as an option to define a concise set of omics features (a so called “molecular signature”), that can describe the most explanatory feature variables (Lê Cao, 2011).

### 2.5 Multi-omics ver3cal integration

*Omix* offers advanced algorithms for sample-level multi-omics integration, designed to accommodate various research scenarios. MOFA and its temporal extension MEFISTO provide a lower-dimensional representation of data by inferring a common latent space, enabling correlations between factors and biologically relevant covariates (Argelaguet et al., 2020). iClusterBayes (Mo et al., 2018) is an effective tool for sample stratification. DIABLO and its continuous counterparts, such as sparse multi-block PLS from mixOmics, facilitate variable selection and biomarker panel identification (Singh et al., 2019). To ensure standardised integrative results across different algorithms, *Omix* employs the custom integrative_results() function. This function extracts multi-omics signatures from the model results, using a custom strategy tailored to the specific model employed. In MOFA, the signature comprises features with the highest weights from a factor strongly correlated with clinical variables of interest. For multiblock PLS, the signature relies on features with the highest set of loadings. In the case of multi-omics clustering, the signature consists of differentially expressed features between clusters. These signatures serve as inputs for downstream analyses, including multiomics networks, module analysis, as well as functional, cell-type, and transcription factor-target enrichment tests. *Omix* also allows users to query external software and databases such as OpenTargets (Ochoa et al., 2021) or Enrichr (Chen et al., 2013) using built-in functions.

## 3 Usage

### 3.1 Pipeline

The core *Omix* pipeline is organised around wrapper functions that receive and return a MultiassayExperiment object. Relevant results are appended to the object’s metadata, enabling users to perform multiple parallel integrative analyses and store the structured integration results within the same object. In terms of model choice, iCluster can be used for sample stratification, DIABLO/MBPLS for biomarker discovery, and MOFA/MEIFESTO for unsupervised integration to explore molecular mechanisms. To demonstrate the comprehensive workflow of an *Omix* analysis, we provide an example that involves two rounds of integration using two distinct algorithms: MOFA and sparse multiblock PLS. This object-oriented structure, combined with wrapper functions greatly enhance the user-friendliness of the analysis, reducing it from thousands of lines of code to approximately 10 (see code implementation in Fig 1).

### 3.2 Pipeline outputs

*Omix*’s core functions operate on the MultiAssay object, taking it as input and producing processed omics data, analysis outputs, and a log of selected parameters, all organised within the same object’s structured metadata slots. This approach offers users the flexibility to directly explore the object’s contents, while also providing the option to generate interactive HTML reports using *Omix*’s pre-designed templates. These include a quality control report, a single omic report, and a multi-omics integration report, which display publication-quality plots, tables, and a comprehensive record of the chosen parameters. For reference, sample reports are accessible on the *Omix* website.

### 3.3 Application

We detailed the use of *Omix* in the Supplementary Material by a case study using Alzheimer’s Disease (AD) bulk proteomics and transcriptomics datasets. We used *Omix* to generate an integrated pseudo-temporal multi-omics profile of AD pathology. We employed the nonsupervised integration method MOFA to project post-mortem samples from different brain regions into a latent space, revealing non-linear inference of AD pathology progression along shared transcriptomics and proteomics variation. This approach identified glial cell activation as a known mechanism of AD pathology progression. Additionally, integrated analysis of proteomics and gene expression uncovered disease progression-specific transcription factor regulatory mechanisms.

## 4 Conclusions

*Omix* is an R package that can be applied in an agnostic manner to integrate bulk transcriptomics and untargeted proteomics datasets. By offering a user-friendly and versatile solution for multi-omics analyses, *Omix* empowers both experts and non-experts to harness the full potential of multi-modal data, in a reproducible fashion, thereby enabling rapid advances in integrative multi-omics research.

## Acknowledgments

We acknowledge the donors and their families for providing human brain 8ssue. Tissue samples were obtained from the London Neurodegenera8ve Diseases Brain Bank at King’s College London, funded by the UK Medical Research Council and the Brains for Demen8a Research programme. Addi8onal 8ssue was provided by the Newcastle Brain Tissue Resource, funded by the UK Medical Research Council, NIHR Newcastle Biomedical Research Centre and Unit, and the Brains for Demen8a Research Programme. Parkinson’s UK Brain Bank at Imperial, funded by Parkinson’s UK, provided samples and data. We thank Diana P. Benitez for her assistance with 8ssue management. ES was supported by the James and Gloria Borley Studentship fund from Imperial College London. PMM acknowledges support from the Edmond J Safra Founda8on, Lily Safra, and an NIHR Senior Inves8gator Award. This work received funding from the UK Demen8a Research Ins8tute, supported by UK DRI Ltd., the UK Medical Research Council, Alzheimer’s Society, and Alzheimer’s Research UK. The study is part of the UK Demen8a Ins8tute Mul8-omics Atlas Project for Alzheimer’s Disease (MAP-AD; map-ad.org).

