## Supplementary material for "Omix: A Multi-Omics Integration Pipeline"

Case study: pseudo-temporal multi-omics profile of Alzheimer's Disease

17 July, 2023

**Step one - Generate a MultiAssayExperiment object** 12

**Step two - Process transcriptomics data** 13

**Step three - Process proteomics data** 14

**Step four - Vertical integration** 14

#### Case study

We illustrate a use case for Omix on a multi-omics dataset comprising bulk proteomics and bulk transcriptomics profiles obtained from 19 Alzheimer's disease (AD) patients and 7 healthy controls. Each sample consisted of both transcriptomic and proteomic datasets. Among the patients, post-mortem samples were collected from two distinct brain regions: the sensorimotor cortex (SOM, n=16), and the middle frontal gyrus (MTG, n=24), which has a higher load of pathology with the disease (Braak & Braak, 1991) (Fig 1). For the majority of patients (14/26), we used multi-omics samples coming from both brain regions, while for the remaining subjects, multi-omics samples were available from either SOM or MTG (2 patients and 10, respectively). All the data can be accessed via a public repository on synapse.org (https://doi.org/10.7303/syn51516099)

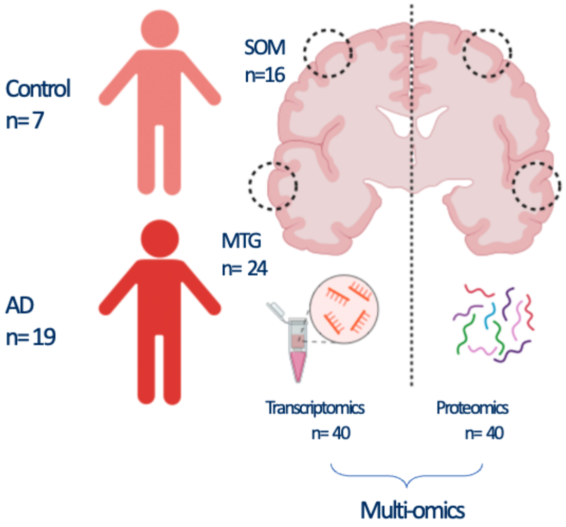

Figure 1: Multi-omics data used for integration

Utilising the integration of these two brain regions, we employ Omix to create a pseudo-temporal multi-omics profile of AD. Put simply, the pseudo-temporal mapping aligns with an increasing pathology load, which is equivalent to a "longer" time spent in disease state. This approach enables us to delve into the underlying biological mechanisms that unfold during the advancement of AD pathology. Following data processing, we utilised the MOFA model for vertical integration, enabling the derivation of a low-dimensional representation capturing shared variation across transcriptomics and proteomics layers. Specifically, factors 6 and 8 emerged as suitable candidates for projecting samples in a latent space that reflects a shared level of explained variation across proteomics and transcriptomics, as opposed to factors where the amount of explained variation is disproportionately attributable to a single modality such as in factors 1-4, 7, and 10-15 (Fig 2A). Second, chosen factors must reflect the variation attributable to the pathological load and/or brain region differences. Factor 6 exhibited significant correlations with the load of tauopathy features such as Braak and phospho-tau (AT8, PHF1), while factor 8 demonstrated correlations with brain regions in the order of when they are affected in disease, thus altogether serving as a proxy for pathology progression (Fig 2B). Altogether, these two factors represent shared variation that is driven by AD pathology at the transcriptomic-proteomic level.

By projecting samples into the integrated latent space, we are able to make a non-linear inference of AD pathology progression along these two axes of variation using the *pseudotime_inference*() function, implemented with Slingshot (Street et al., 2018) (Fig 2C). This inferred pseudotime well depicts AD pathology progression between brain regions. Factor 6, by separating pathological features from anatomical one, provides a common framework across datasets with which one can define the molecular progression of AD pathology. To elucidate the molecular features driving factor 6, we identified top genes and proteins based on their weights and grouped them into co-expression modules (ME1-5) using Omix's *extract_weights*() and *multiomics_modules*() functions. Notably, modules 1, 3 and 4 displayed strong correlation of their eigenvalues with known neuropathological hallmarks of AD (Fig 2D). The convergence of multiomics modules and pseudo-time analysis provides a way to estimate synchronised protein and mRNA changes across the disease trajectory, shedding light on the relative magnitudes of these changes during AD pathology progression (Fig 2E). Indeed, we found that module expression changes along the inferred trajectory also describes related glial pathology related to microglial and astrocyte activation characterised via pathway enrichment in ME4 and ME1, respectively. These were also significantly correlated to the inferred pseudotime (Fig 2D) and displayed reduced levels in the early-stage region, SOM, compared to MTG ($p$ value $< 0.05$) (Fig 3) . The integrated transcriptomic-proteomic network analysis was used to infer on regulatory mechanisms. For example, we identified two transcription factors CEBPB and STAT3 in the proteomics, for which the target genes were significantly enriched in the astrocyte (ME1) and microglia (ME4) modules respectively (Fig 4), consistent with previous studies highlighting their role in regulating a neuroinflammatory response in AD (Millot et al., 2020, Yao et al., 2021 ). This reinforces the power of using multi-omics pseudo-temporal approach to study regulatory mechanisms of progression. More so, module features could serve as potential novel therapeutic targets or progression biomarkers. Finally, this workflow could be replicated for factors 1, 10, and 15, which also display significant correlations to neuropathological hallmarks though depicting a single modality level of variation, to highlight modality differences and provide a different analysis angle under the same pathological conditions.

Results of this case study underscore the promising prospects of utilising multi-omics integration to enhance our comprehension of molecular processes implicated in diseases such as AD. We illustrated an application of the capacity of this approach to uncover established mechanisms of AD progression by employing transcriptomic-proteomic integration and pseudo-time inference.

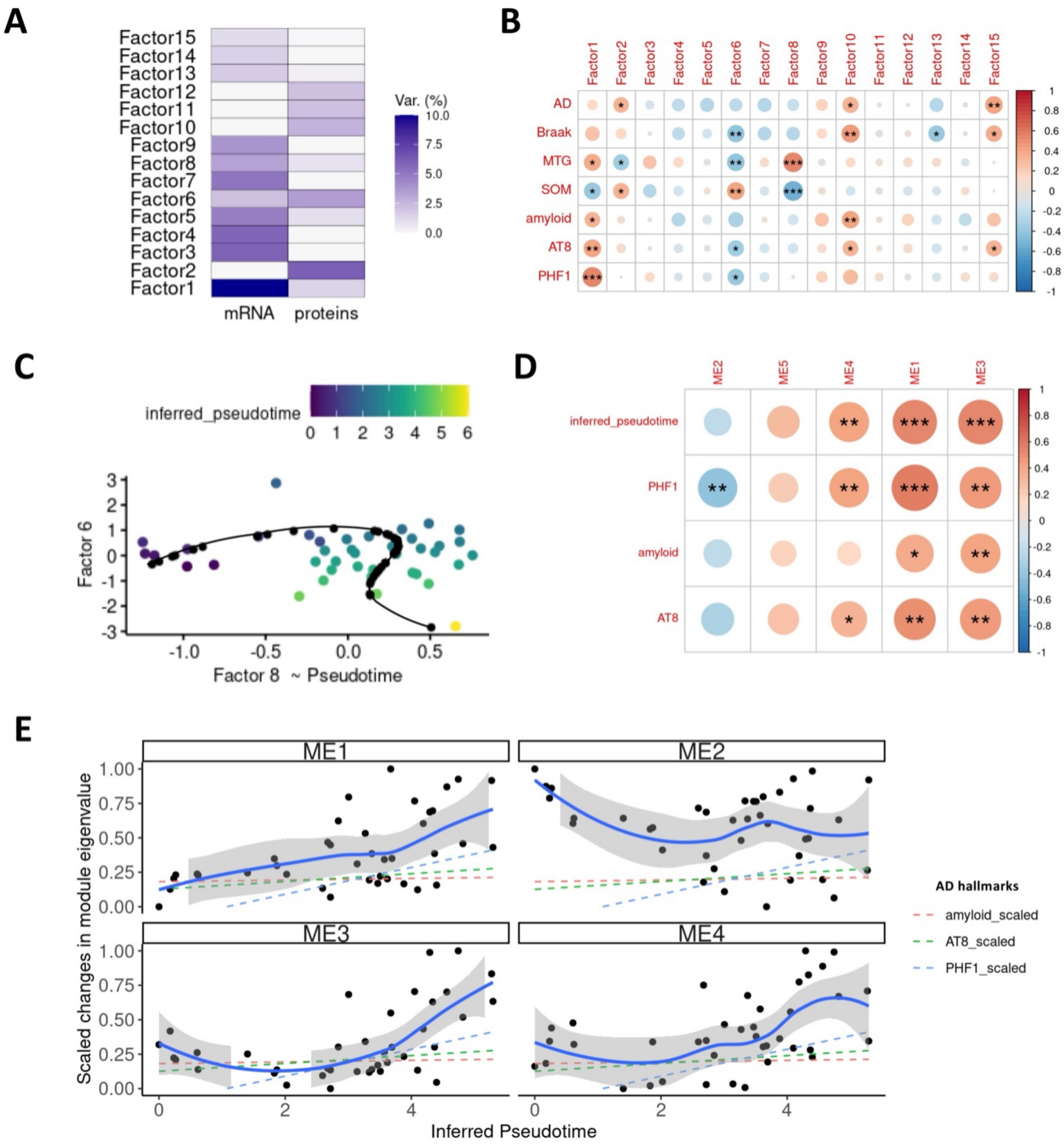

Figure 2: Pseudo-temporal multi-omics profile of Alzheimer's disease. Panel (A) Displays the variance explained by the learnt factors across transcriptomics and proteomics. Panel (B) displays a correlation heatmap depicting the relationship between MOFA-derived multi-omics factors and neuropathological covariates, quantified using Pearson's coefficient. The size of each dot corresponds to the absolute correlation coefficient. MTG and SOM were one hot encoded based on sample regional provenance and Pearson's correlation reflects the Point-Biserial Correlation here (C), Slingshot-based pseudotime inference is employed utilising embeddings derived from factors 6 and 8. Samples are projected into a latent space, with colours indicating the inferred trajectory (depicted by the black curve). Panel (D) presents a correlation heatmap showcasing the relationship between multi-omics modules and AD neuropathological hallmarks, evaluated using Pearson's correlation coefficient. To assess changes in module eigenvalues across pseudotime, Panel (E) exhibits scaled changes accompanied by smoothed Loess regression curves, with the grey banner representing the standard error. Additionally, a linear fit illustrates the scaled changes of AD hallmarks. *p<0.05, **p<0.01, ***p<0.001.

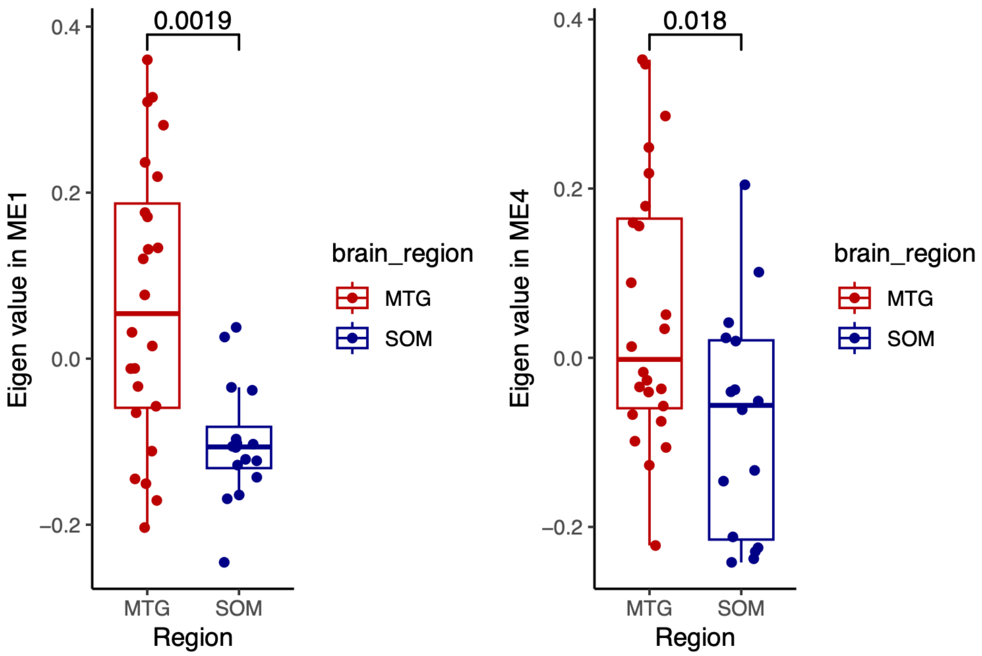

Figure 3: Glial modules expression in MTG and SOM. Eigen values of each module represent the first principal component of the module expression. ME1 is the astrocyte module and ME4 is the microglia module. Brain region differences are assessed using Wilcoxon rank-sum tests.

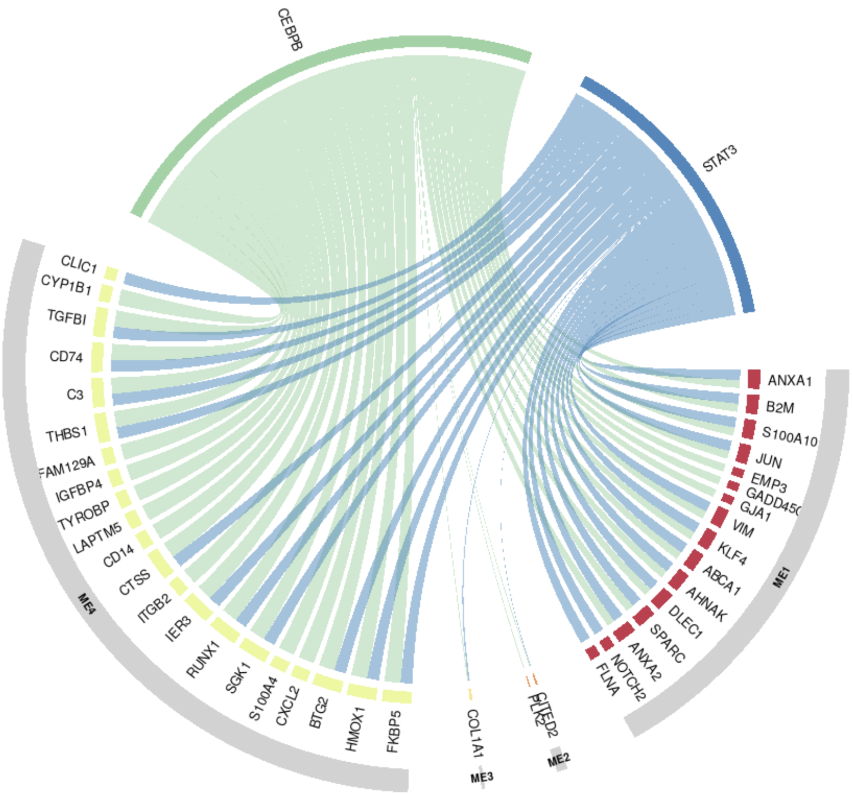

Figure 4: Transcription Factor to target genes relationship. An integrated network analysis was employed to gain insights into the regulatory mechanisms involved in AD progression. We used an established Transcription Factor (TF)-target gene library from ChIP-seq data to identify target genes enriched in the progression modules (ME1-4), based on TF identified from protein abundance. The circos plot visualises the relationship between two identified transcription factors, CEBPB and STAT3, and their associated gene targets, which exhibited co-expression patterns during AD progression.

### Reproducing The Analysis

The goal of Omix is to provide tools in R to build a complete analysis workflow for integrative analysis of data generated from multi-omics platform.

- Generate a multi-omics object using `MultiAssayExperiment`.
- Quality control of single-omics data.
- Formatting, normalisation, denoising of single-omics data.
- Separate single-omics analyses.
- Integration of multi-omics data for combined analysis.
- Publication quality plots and interactive analysis reports based of shinyApp.

Currently, Omix supports the integration of bulk transcriptomics and bulk proteomics.

#### Running *Omix*

The `Omix` pipeline requires the following input:

- `rawdata_rna`: A data-frame containing raw RNA count with rownames as gene and colnames as samples.
- `rawdata_protein`: A data-frame containing protein abundance with rownames as proteins and colnames as samples.
- `map_rna`: A data-frame of two columns named `primary` and `colname` where primary should contain unique sample name with a link to sample metadata and colname is the column names of the `rawdata_rna` data-frame.
- `map_protein`: A data-frame of two columns named `primary` and `colname` where primary should contain unique sample name with a link to sample metadata and colname is the column names of the `rawdata_protein` data-frame.
- `metadata_rna`: A data-frame containing `rna` assay specific metadata where rownames are same as the colnames of `rawdata_rna` data.frame.
- `metadata_protein`: A data-frame containing `protein` assay specific metadata where rownames are same as the colnames of `rawdata_protein` data.frame.
- `individual_metadata`: A data-frame containing individual level metadata for both omics assays.

Call required packages for vignette:

```
library(Omix)
library(purrr)
library(ggplot2)
library(dplyr)
library(synapser)
library(tibble)
```

First we must load the data from Omix for the vignette:

All the data is stored on synapse: https://doi.org/10.7303/syn51516099

```
outputDir <- tempdir()
ctd_fp <-file.path(outputDir, "ctd.rds")
ensembl_fp  <-file.path(outputDir, "ensembl.rds")
TF_fp <- file.path(outputDir, "Enrichr_Queries.gmt.txt")
GO_fp <- file.path(outputDir, "GO_Biological_Process_2021.txt")

path <- "https://raw.githubusercontent.com/eleonore-schneeg/OmixData/main/"
```

```r
download.file(url = paste0(path,"ctd-2.rds"),
              destfile=ctd_fp ,
              headers = c( Authorization=paste("token", Sys.getenv("GH_TOKEN"))))

download.file(url = paste0(path,"ensembl_mappings_human.tsv"),
              destfile=ensembl_fp ,
              headers = c(Authorization = paste("token", Sys.getenv("GH_TOKEN"))))

download.file(url = paste0(path,"Enrichr_Queries.gmt.txt"),
              destfile=TF_fp ,
              headers = c(Authorization = paste("token", Sys.getenv("GH_TOKEN"))))

download.file(url = paste0(path,"GO_Biological_Process_2021.txt"),
              destfile=GO_fp ,
              headers = c(Authorization = paste("token", Sys.getenv("GH_TOKEN"))))

synapser::synLogin(email=Sys.getenv("SYNAPSE_ID"), password=Sys.getenv("SYNAPSE_PASSWORD"))
```

Welcome, ems2817!NULL

```r
rawdata_rna <- read.csv(synapser::synGet('syn51516100')$path,
                          header=T,
                          stringsAsFactors = F,
                          row.names=1)

rawdata_protein <- read.csv(synapser::synGet('syn51516105')$path,
                              header=T,
                              stringsAsFactors = F,
                              row.names=1)

map_rna <-read.csv(synapser::synGet('syn51516102')$path,
                      header=T,
                      stringsAsFactors = F,
                      row.names=1)

map_prot <- read.csv(synapser::synGet('syn51516104')$path,
                       header=T,
                       stringsAsFactors = F,
                       row.names=1)

metadata_rna<-read.csv(synapser::synGet('syn51516103')$path,
                           header=T,
                           stringsAsFactors = F,
                           row.names=1)

metadata_prot <- read.csv(synapser::synGet('syn51516106')$path,
                            header=T,
                            stringsAsFactors = F,
                            row.names=1)

individual_metadata <-read.csv(synapser::synGet('syn51516101')$path,
                                  header=T,
                                  stringsAsFactors = F,
                                  row.names=1)
```

```r
clusters<-readRDS(synapser::synGet('syn51516107')$path)
integrative_model<-readRDS(synapser::synGet('syn51520491')$path)

ctd <-  readRDS(paste0(outputDir,'/ctd.rds'))
ensembl <-read.delim(file=paste0(outputDir,'/ensembl.rds'),
                        sep = '\t',
                        header = TRUE)

rna_qc_data_matrix <- NULL
```

Sanity check

```r
all(rownames(metadata_rna) == colnames(rawdata_rna))
```

```
## [1] TRUE
```

```r
all(rownames(metadata_prot) == colnames(rawdata_protein))
```

```
## [1] TRUE
```

##### Raw rna counts

`rawdata_rna` is a data-frame of raw counts, with features as rows and samples as columns

```r
print(rawdata_rna[1:5,1:5])
```

```
##                 IGF117745 IGF117748 IGF117756 IGF117758 IGF117759
## ENSG00000223972         0         0         1         0         1
## ENSG00000227232        23        26        40        26        30
## ENSG00000278267         7         5         1         4         0
## ENSG00000243485         0         0         0         1         0
## ENSG00000284332         0         0         0         0         0
```

##### Raw protein abundance

`rawdata_protein` is a data-frame of raw protein abundances, with features as rows and samples as columns

```r
print(rawdata_protein[10:15,10:15])
```

```
##                    BBN_16213_SOM_P2WH4_001 BBN_18399_MTG_P2WF5_001
## MAP1LC3B;MAP1LC3B2               140.21600                118.1090
## PGP                               33.91880                 24.3759
## BTBD17                                  NA                      NA
## WIPF3                             39.10300                 98.0357
## IGLON5                             8.55527                 10.2548
## TUBAL3                          9432.14000              10277.5000
##                    BBN_18399_SOM_P2WF6_001 BBN_18813_SOM_P3WB2_001
## MAP1LC3B;MAP1LC3B2                116.8410               109.55500
## PGP                                15.2674                31.62250
## BTBD17                             13.8635                      NA
## WIPF3                              68.9909                26.42480
## IGLON5                             21.7657                 8.52512
## TUBAL3                           9565.3200             10144.50000
##                    BBN_19214_MTG_P2WA3_001 BBN_19214_SOM_P2WA4_001
## MAP1LC3B;MAP1LC3B2                121.1260                166.2360
## PGP                                18.2978                 26.0274
## BTBD17                                  NA                 17.6870
```

```
## WIPF3                              58.8438                  NA
## IGLON5                             12.5892             20.8584
## TUBAL3                           9069.5600          10454.6000
```

##### Maps

Maps are data frame containing two columns: `primary` and `colname`. The `primary` column should be the individual ID the individual metadata, and the `colname` the matched sample names from the raw matrices columns. If sample and individual ids are the same, maps aren't needed (primary and colnames are the same).

```
print(head(map_prot))
```

```
##         primary                 colname
## 1 BBN_10099-MTG BBN_10099_MTG_P3WH10_001
## 2 BBN_10099-SOM  BBN_10099_SOM_P4WA1_001
## 3 BBN_10109-MTG  BBN_10109_MTG_P4WA3_001
## 4 BBN_10109-SOM  BBN_10109_SOM_P4WA4_001
## 5 BBN_10611-MTG BBN_10611_MTG_P2WD10_001
## 6 BBN_10611-SOM  BBN_10611_SOM_P2WE1_001
```

```
print(head(map_rna))
```

```
##             primary   colname
## 1  BBN002.30035-SOM IGF117745
## 4      BBN_7519-SOM IGF117748
## 12 BBN003.32235-MTG IGF117756
## 14    BBN_21792-SOM IGF117758
## 15     BBN_7617-MTG IGF117759
## 16     BBN_9984-SOM IGF117760
```

##### Technical metadata

Technical metadata are data-frames contain the column `colname` that should match the sample names in the raw matrices, and any additional columns related to technical artefacts like batches

```
print(head(metadata_rna))
```

```
##                  sample_id sample_name RIN Amyloid_Beta       pTau
## IGF117745 BBN002.30035-SOM   IGF117745 6.4     0.343451 0.00553998
## IGF117748     BBN_7519-SOM   IGF117748 8.1           NA         NA
## IGF117756 BBN003.32235-MTG   IGF117756 2.7     2.027670 6.98931000
## IGF117758    BBN_21792-SOM   IGF117758 5.3     1.529590 0.04699940
## IGF117759     BBN_7617-MTG   IGF117759 2.0     3.262270 2.55134000
## IGF117760     BBN_9984-SOM   IGF117760 7.7     3.653590 1.65894000
##           pct_pTau_Positive       PHF1
## IGF117745         0.0426309 0.00121131
## IGF117748                NA         NA
## IGF117756         5.2597500 2.42065000
## IGF117758         0.1894760 0.00342135
## IGF117759         0.3749760 1.56408000
## IGF117760         1.5665900 0.06801730
```

```
print(head(metadata_prot))
```

```
##                              sample_id              sample_name batch
## BBN_10099_MTG_P3WH10_001 BBN_10099-MTG BBN_10099_MTG_P3WH10_001    P3
## BBN_10099_SOM_P4WA1_001  BBN_10099-SOM  BBN_10099_SOM_P4WA1_001    P4
## BBN_10109_MTG_P4WA3_001  BBN_10109-MTG  BBN_10109_MTG_P4WA3_001    P4
```

```
## BBN_10109_SOM_P4WA4_001  BBN_10109-SOM  BBN_10109_SOM_P4WA4_001    P4
## BBN_10611_MTG_P2WD10_001 BBN_10611-MTG BBN_10611_MTG_P2WD10_001    P2
## BBN_10611_SOM_P2WE1_001  BBN_10611-SOM  BBN_10611_SOM_P2WE1_001    P2
```

##### Individual metadata

`individual_metadata` should contain individual level metadata, where one column matches the `primary` column in maps.

```
print(head(individual_metadata))
```

```
##       sample_id brain_region age    BBN_ID sex diagnosis apoe_final Braak PMD
## 1 BBN_10099-MTG          MTG  91 BBN_10099   F   Control      E3/E3     2  22
## 2 BBN_10099-SOM          SOM  91 BBN_10099   F   Control      E3/E3     2  22
## 3 BBN_10109-MTG          MTG  80 BBN_10109   F   Control      E2/E3     2  23
## 4 BBN_10109-SOM          SOM  80 BBN_10109   F   Control      E2/E3     2  23
## 5 BBN_10611-MTG          MTG  92 BBN_10611   F        AD      E3/E3     3  30
## 6 BBN_10611-SOM          SOM  92 BBN_10611   F        AD      E3/E3     3  30
##    amyloid       pTau       PHF1 AD MTG SOM
## 1 0.585285 0.00649899 0.00168676  0   1   0
## 2 0.226991 0.00960187         NA  0   0   1
## 3 0.782684 0.92852500 0.67876900  0   1   0
## 4 0.059688 0.00193356         NA  0   0   1
## 5 0.307882 0.01312750 0.01797600  1   1   0
## 6 0.530825 0.00562307 0.00824549  1   0   1
```

##### Optional inputs

Optional inputs contain additional data frame used for QC visualisation purposes

```
print(head(rna_qc_data_matrix))
```

```
## NULL
```

#### Step one - Generate a MultiAssayExperiment object

To run Omix, we first need to generate a multi-omics object The package currently supports transcriptomics and proteomics bulk data only.

```
individual_metadata$diagnosis <- factor(individual_metadata$diagnosis,
                                        levels = c("Control", "AD"))
individual_metadata$sex <- factor(individual_metadata$sex)

multiomics_object=generate_multiassay(rawdata_rna =rawdata_rna,
                                    rawdata_protein = rawdata_protein,
                                    individual_to_sample=FALSE,
                                    map_rna = map_rna,
                                    map_protein = map_prot,
                                    metadata_rna = metadata_rna,
                                    metadata_protein = metadata_prot,
                                    individual_metadata = individual_metadata,
                                    map_by_column = 'sample_id',
                                    rna_qc_data=FALSE,
                                    rna_qc_data_matrix=NULL,
                                    organism='human')
```

```
## v Ensembl ID conversion to gene symbol

## v Retrieval of gene biotype

## v RNA raw data loaded

## class: SummarizedExperiment
## dim: 58884 48
## metadata(1): metadata
## assays(1): rna_raw
## rownames(58884): ENSG00000223972 ENSG00000227232 ... ENSG00000277475
##   ENSG00000268674
## rowData names(3): ensembl_gene_id gene_name gene_biotype
## colnames(48): IGF117745 IGF117748 ... IGF117836 IGF117837
## colData names(7): sample_id sample_name ... pct_pTau_Positive PHF1

## v Protein raw data loaded

## class: SummarizedExperiment
## dim: 3228 82
## metadata(1): metadata
## assays(1): protein_raw
## rownames(3228):
##   IGKV2-28;IGKV2-29;IGKV2-30;IGKV2-40;IGKV2D-26;IGKV2D-28;IGKV2D-29;IGKV2D-30;IGKV2D-40
##   IGKV3-11;IGKV3D-11 ... FAM169A SEC23IP
## rowData names(1): gene_name
## colnames(82): BBN_10099_MTG_P3WH10_001 BBN_10099_SOM_P4WA1_001 ...
##   BBN003.35520_MTG_P2WD4_001 BBN003.35521_SOM_P2WD5_001
## colData names(3): sample_id sample_name batch

## v MultiAssayExperiment object generated!
```

The MultiAssayExperiment object was succesfully created. The following steps will process and perform QC on each omic layers of the `multiomics_object` object.

#### Step two - Process transcriptomics data

```
multiomics_object=process_rna(multiassay=multiomics_object,
            transformation='rlog',
            protein_coding=TRUE,
            min_count = 10,
            min_sample = 0.5,
            dependent =  "diagnosis",
            levels = c("Control","AD"),
            covariates=c('age','sex','PMD'),
            filter=TRUE,
            batch_correction=TRUE,
            batch=NULL,
            remove_sample_outliers= FALSE)
```

```
## v NORMALISATION & TRANSFORMATION

## v GENE FILTERING

## v Keeping only protein coding genes

## v 39155 / 58884 non protein coding genes were dropped

## v 19729 protein coding genes kept for analysis
```

```
## v QC parameters selected require genes to have at least 50 % of samples with counts of 10 or more
## v 4849 / 19729 genes were dropped
## v 14880 genes kept for analysis
## v RLOG TRANSFORMATION
## v BATCH CORRECTION
## v Using Limma on  to remove technical artefacts, and age as biological confoundersUsing Limma on  to
## v Transcriptomics data processed!
## v Processing parameters saved in metadata
```

#### Step three - Process proteomics data

```
multiomics_object=process_protein(
    multiassay=multiomics_object,
    filter=TRUE,
    min_sample = 0.5,
    dependent =  "diagnosis",
    levels = c("Control","AD"),
    imputation = 'minimum_value',
    remove_feature_outliers= FALSE,
    batch_correction= FALSE,
    batch="batch",
    correction_method="median_centering",
    remove_sample_outliers=FALSE,
    denoise=TRUE,
    covariates=c('PMD','sex','age'))
```

```
## v SCALING NORMALIZATION
## v FILTERING
## v 283 / 3228 proteins filtered
## v 2945 proteins kept for analysis
## v IMPUTATION
## v Imputation of remaining missing values based on
## 50% of minimum level of abundance for each protein
## v DENOISING BIOLOGICAL COVARIATES
## v Using Limma on PMD as biological confoundersUsing Limma on sex as biological confoundersUsing Limma
```

#### Step four - Vertical integration

**Omix** supports a range of vertical integration models:

- Possible integration methods are `MOFA`,`DIABLO`,`sMBPLS`,`iCluster`,`MEIFESTO`
- The choice of the integration models depends on the research use case of interest.
- Here we display the use of **Omix** on a popular integration method, MOFA or Multi-Omics Factor analysis (Argelaguet et al. 2018).

In this vignette, we proceed with a pseudotemporal multi-omics integration of 40 brain Alzheimer's disease (AD) and Control samples coming from two brain regions

- The somatosensory cortext (SOM)
- The middle frontal gyrus (MTG)

The MTG is known to be affected earlier during AD progression, while SOM at later stages. Using these two regions as pseudotemporal proxi in the integrative process, we are able to gain a deeper understanding of biological mechanisms that occur during AD progression.

##### Pseudotemporal multi-omics integration using MOFA

```
multiomics_object=vertical_integration(multiassay=multiomics_object,
                                slots = c(
                                   "rna_processed",
                                   "protein_processed"
                                 ),
                                integration='MOFA',
                                ID_type = "gene_name",
                                dependent='diagnosis',
                                intersect_genes = FALSE,
                                num_factors = 15,
                                scale_views = FALSE,
                                most_variable_feature=TRUE)
```

The integration process is successful and all integration related object are stored in the `integration` slot of the multi-omics object.

```
print(multiomics_object@metadata$integration$MOFA)
```

```
## Trained MOFA with the following characteristics:
##  Number of views: 2
##  Views names: mRNA proteins
##  Number of features (per view): 2945 2945
##  Number of groups: 1
##  Groups names: group1
##  Number of samples (per group): 40
##  Number of factors: 15
```

##### Optional: load a pretrained model

Since package version slightly affect model outputs, we reload a pre-trained model.

```
multiomics_object@metadata$integration$MOFA=integrative_model
```

#### Step five - Post integration downstream analyses

**Omix** provides a range of built-in downstream analyses functions and visualisations. All downstream analyses will be performed on the integrated object stored in the `integrated` slot.

##### Load integrated object

```
integrated_object=multiomics_object@metadata$integration$MOFA
metadata=integrated_object@samples_metadata
```

#### Factor explorations

##### Variance explained

```
MOFA2::plot_variance_explained(integrated_object, max_r2=10)
```

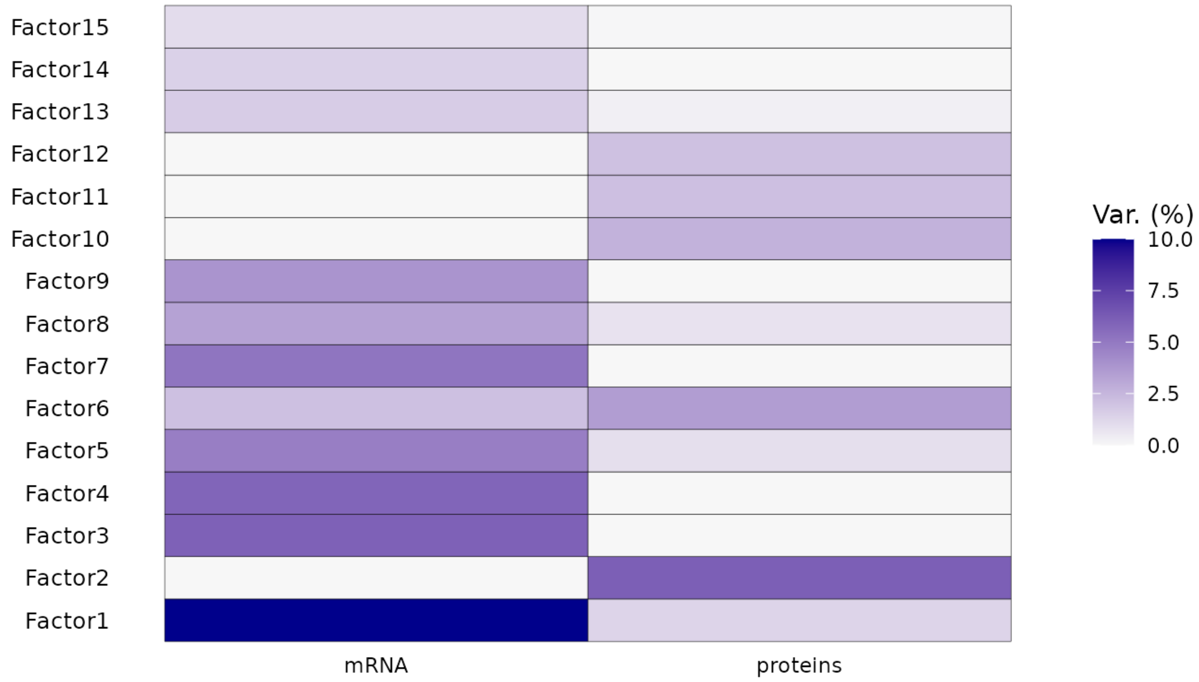

##### Correlation of factors with covariates

```
integrated_object@samples_metadata$SOM=ifelse(integrated_object@samples_metadata$brain_region=='SOM',1,[illegible]
plot=correlation_heatmap(integrated_object,
                    covariates=c("AD","Braak",'MTG','SOM',
                                 'amyloid','pTau','PHF1'))
```

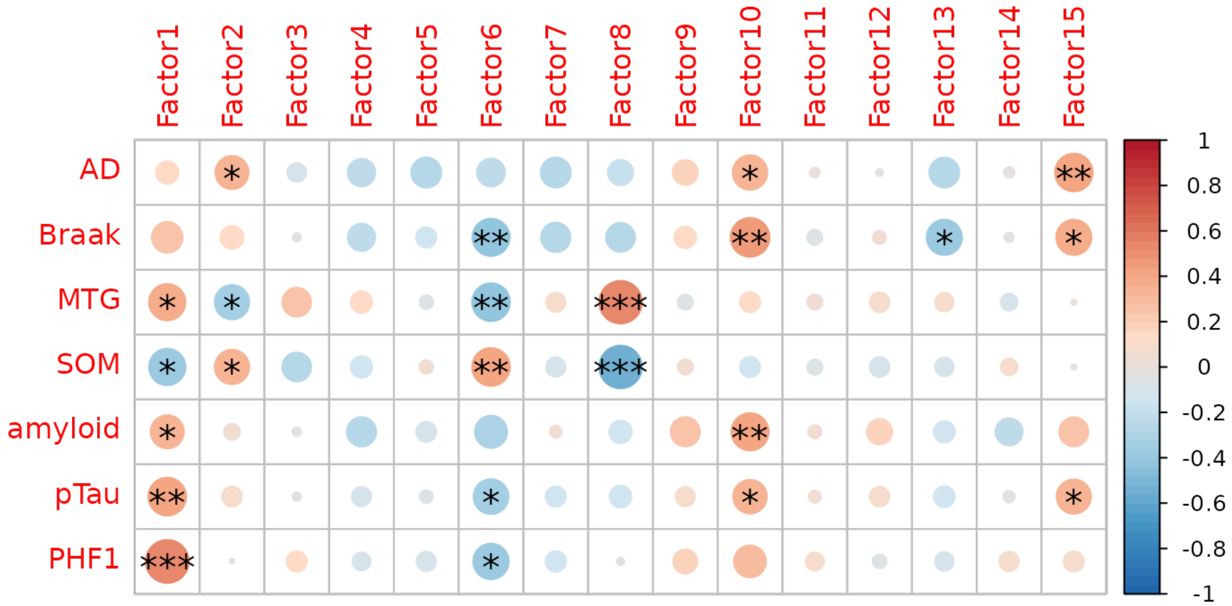

Factors 6 and 8 will be chosen for downstream analysis:

- Factor 6: Is strongly correlated with neuropathological variables including PHF1 and pTau (AT8).
- Factor 8: Is strongly correlated with pseudotemporal covariates, MTG and SOM
- Both share some level of variance explained by the transcriptomics and proteomics layers

  Consequentially, If we project samples on these two axis of variation, we can infer on the pseudotemporal trajectory of neuropathology at the RNA and protein level.

#### Integrated latent space

##### Chosen integrated axes of variation

```
MOFA2::plot_factors(integrated_object,
             factors=c(8,6),
             color_by='PHF1', shape_by = 'brain_region', scale = T)+
              xlab('Factor 8 ~ Pseudotime' )+
              ylab('Factor 6 ~ Neuropathology' )
```

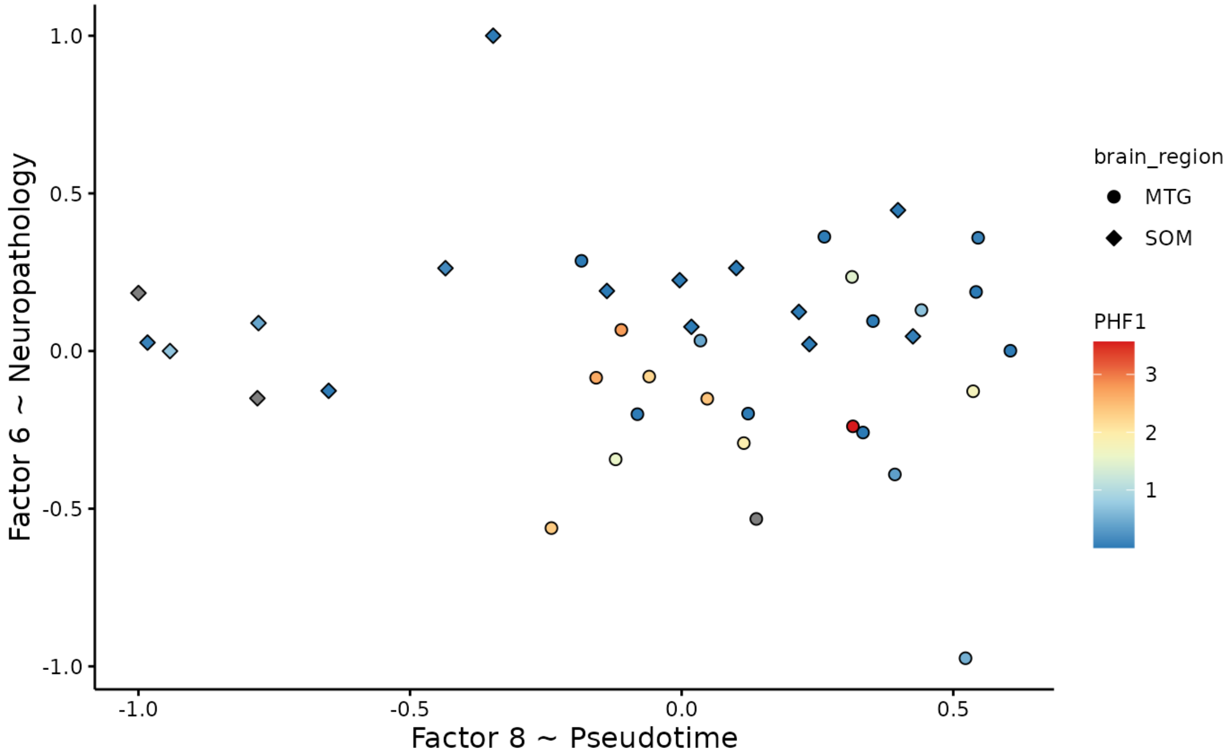

##### Pseudotime inference with Slingshot

The goal of `slingshot` (Street et al. 2017), is to use clusters of cells to uncover global structure and convert this structure into smooth lineages represented by one-dimensional variables, called "pseudotime."

> Here, we extend the use of `slingshot` to infer on individual level pseudotime to infer on Alzheimer's disease trajectory.

`slingshot` takes a matrix of embeddings in reduced dimension as input, which here are Factors 8 and 6 from the `MOFA` multi-omics integration. We can optionally specify the cluster to start or end the trajectory based on biological knowledge. Here, the starting cluster should be `1` as it coincides with lowest neuropathology levels and most "recently" affected region, SOM, in terms of AD trajectory.

```
#clusters=MOFA2::cluster_samples(integrated_object,
#                          k=3,
#                          factors = c(8,6))

#load clusters for reproducibility
MOFA2::plot_factors(integrated_object,
             factors=c(8,6),
             color_by=clusters$cluster, scale = F)
```

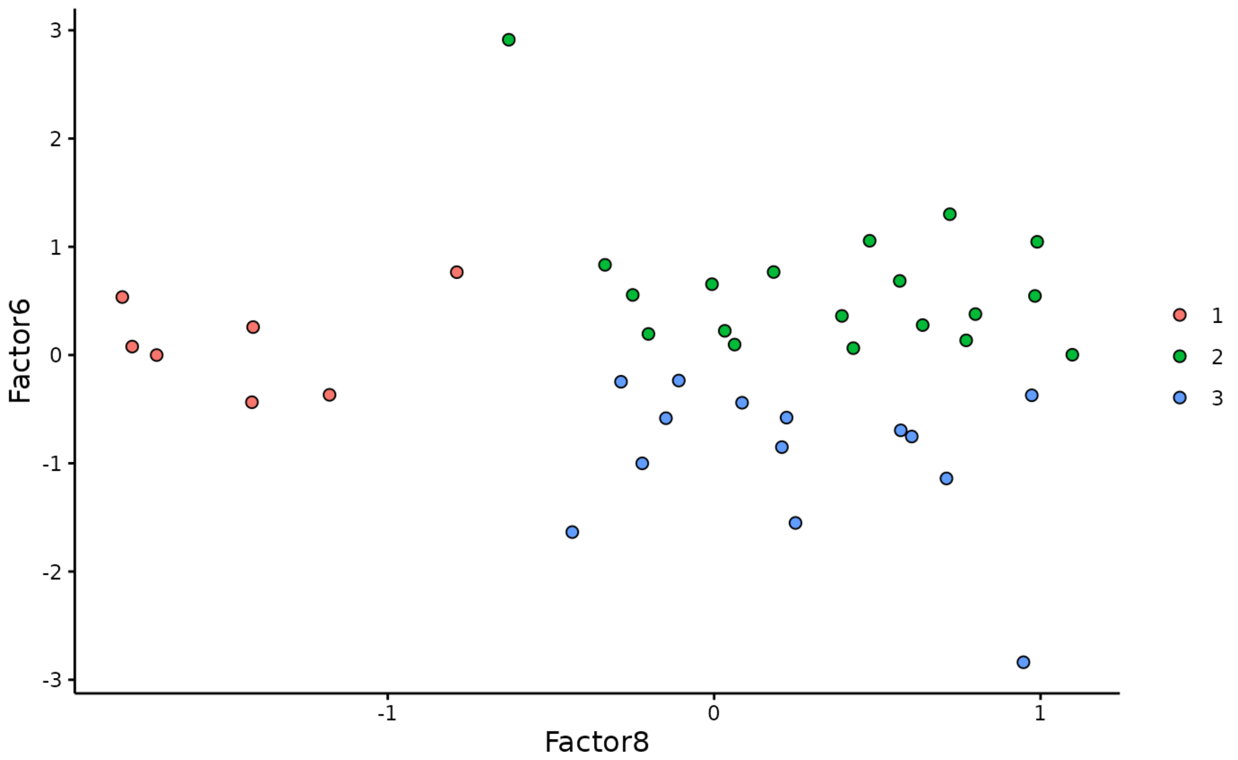

```
pseudotime=pseudotime_inference(integrated_object,
                                 clusters,
                                 time_factor=8,
                                 second_factor=6,
                                 start.clus = 1,
                                 end.clus=3,
                                 lineage = 'Lineage1')

pseudotime$plot
```

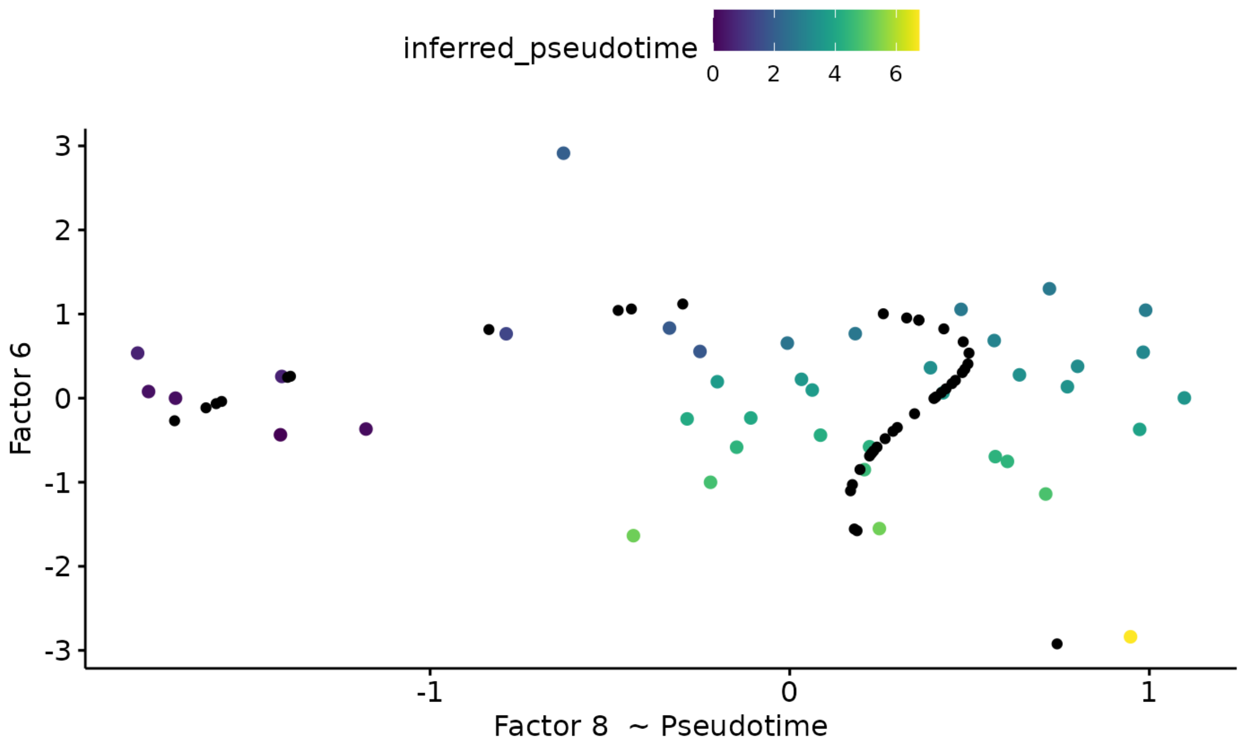

**Update the integrated_object with the pseudotime**

```
multiomics_object@metadata$integration$MOFA=pseudotime$model
integrated_object=multiomics_object@metadata$integration$MOFA
MOFA2::samples_metadata(integrated_object)$inferred_pseudotime
```

```
##  [1] 4.3991398 0.0000000 3.2695341 0.6197033 3.3734278 3.5719823 3.1405570
##  [8] 3.3321288 3.5080573 3.6197082 1.4029378 3.6919371 2.0234374 2.7175514
## [15] 2.5931510 4.3672023 1.8283014 5.2932327 4.7431202 3.4678981 4.3375355
## [22] 0.2645565 4.5626404 0.6050538 1.8689173 2.8424525 2.7175514 4.2916315
## [29] 2.6748828 4.0987023 0.1769606 4.8168531 3.0058323 3.6727096 6.7492822
## [36] 3.8811122 0.2341031 5.2714195 4.1889176 4.0515236
```

#### Extract features that are driving Factor 6 based on feature weights

Weights vary from -1 to +1, and provide a score for how strong each feature relates to each factor, hence allowing a biological interpretation of the latent factors. Features with no association with the factor have values close to zero, while genes with strong association with the factor have large absolute values.

The sign of the weight indicates the direction of the effect:

- A positive weight indicates that the feature has higher levels in the samples with positive factor values.
- A negative weight indicate higher levels in samples with negative factor values.

The `extract_weigths`function enable to extract the weights on the desired factor at a defined absolute threshold, and return an object with positive and negative weights above and below this threshold respectively. It also returns QC plots

- A distribution of the feature weights for each omic layer at the designated factor

- The relationship between the feature/`sense_check_variable` correlation and weights. High weights should coincide with stronger correlation if the `sense_check_variable` is an important driver of variation in the designated factor.

```
weights=extract_weigths(integrated_object,
                        factor=6,
                        threshold=0.2,
                        sense_check_variable='PHF1')
```

###### Weights distribution in Factor 6

```
weights$distribution_plot$rna
```

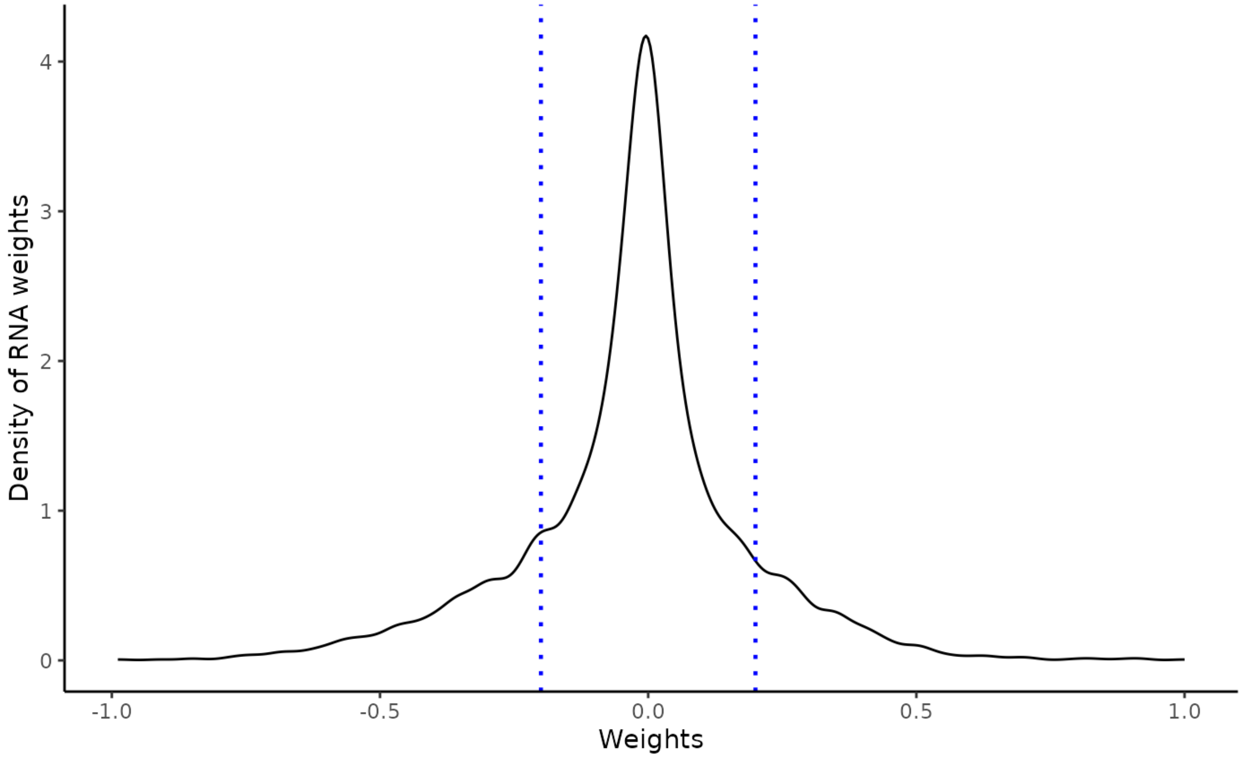

```
weights$distribution_plot$protein
```

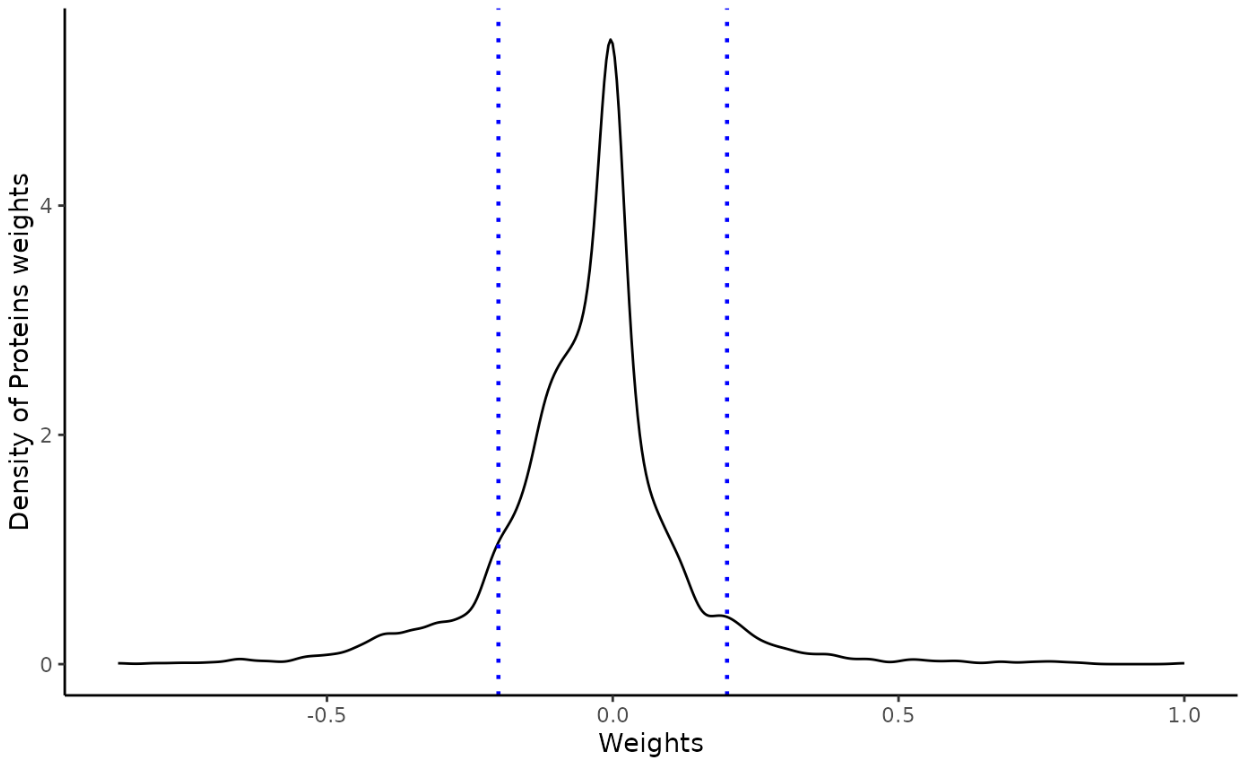

**Correlation to PHF1 and weights in Factor 6**

```
weights$weights_cor_plot$rna
```

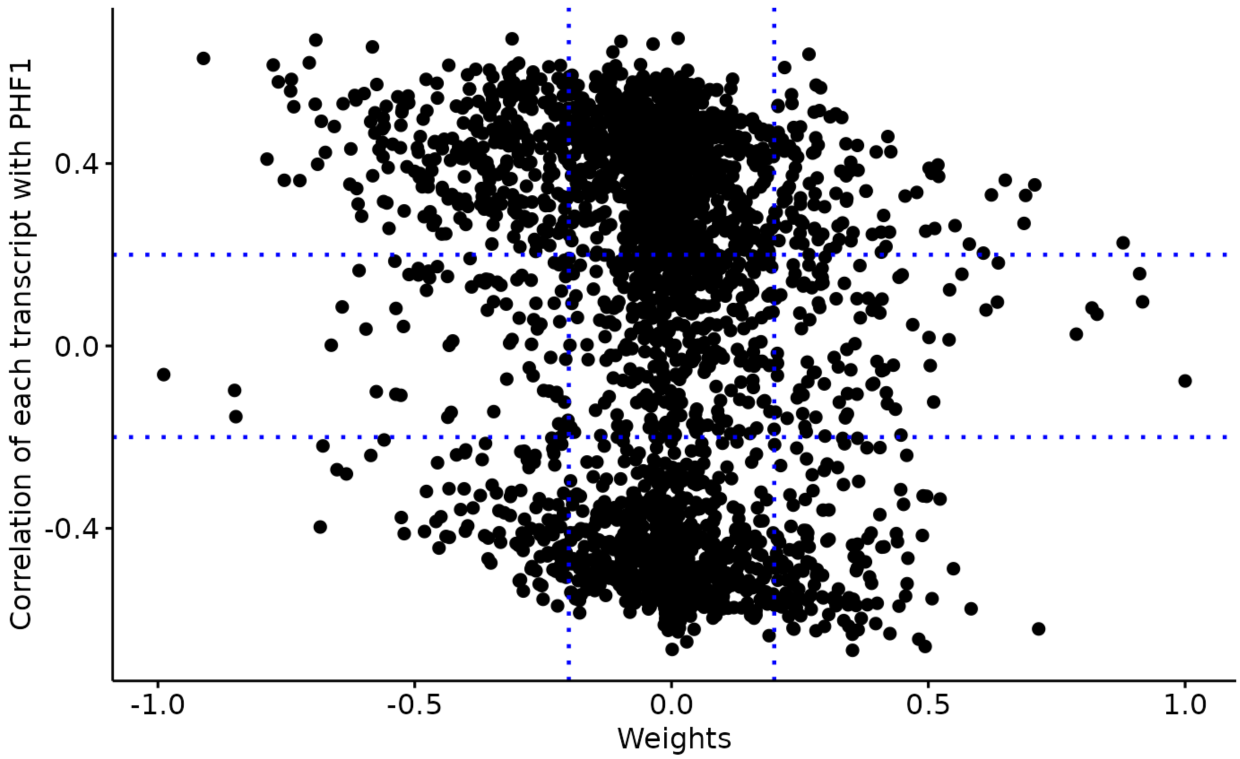

```
weights$weights_cor_plot$protein
```

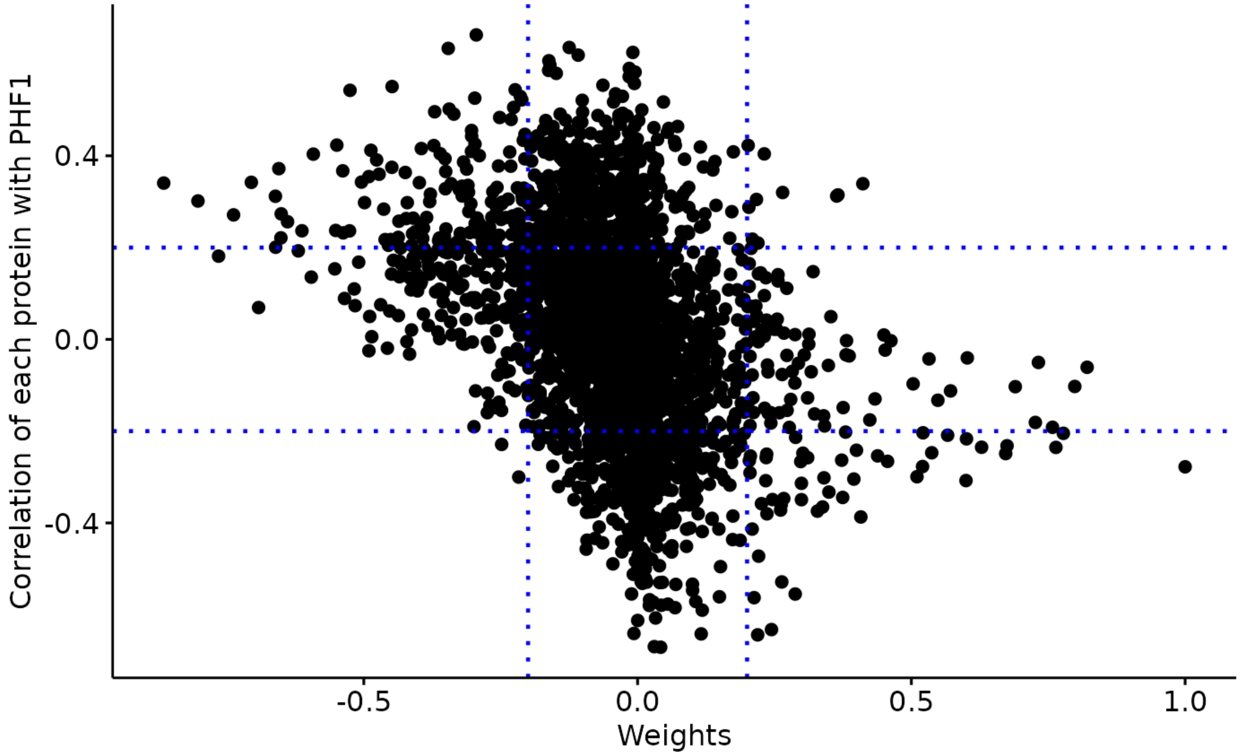

##### Negative weights in factor 6: Up-regulated in later stage AD

A negative weight indicates that the feature has higher levels in the samples with negative factor values, which in this analysis coincides with later Braak stages.

```
integrated_object@samples_metadata$Braak=as.factor(integrated_object@samples_metadata$Braak)
MOFA2::plot_factor(integrated_object,
            factors = 6,
            color_by = "Braak",
            add_violin = TRUE,
            dodge = TRUE
)
```

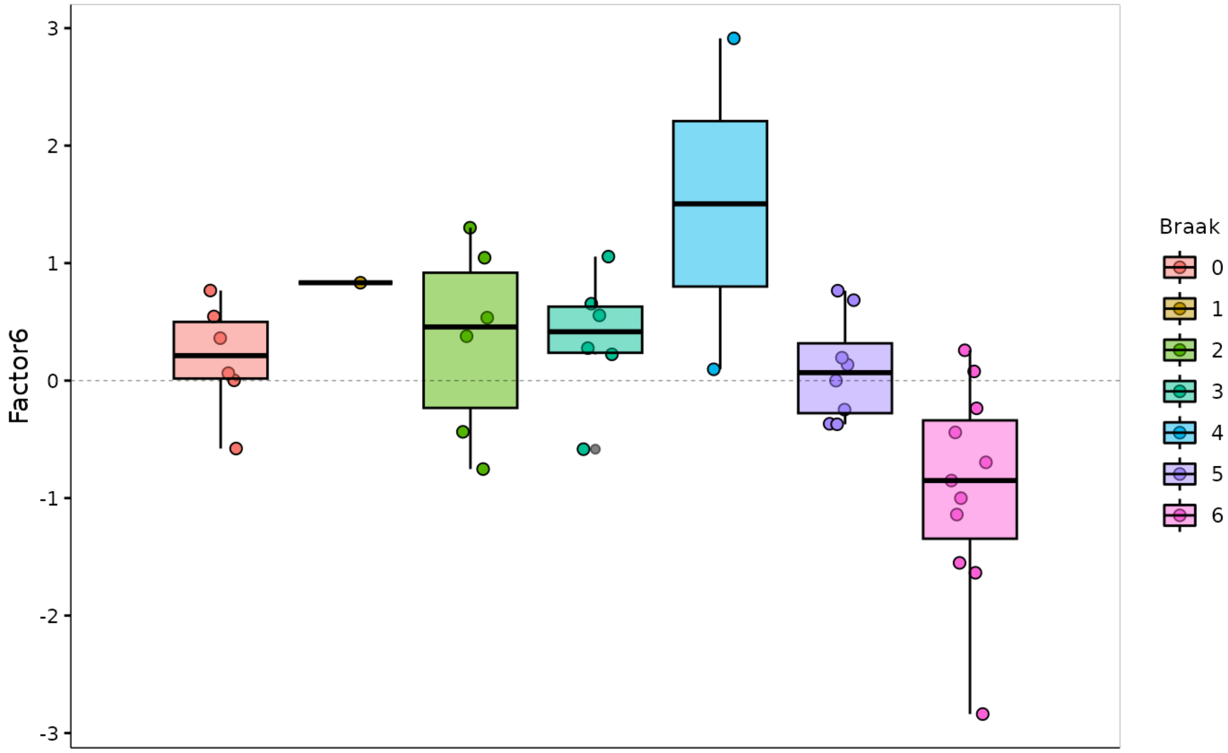

**Selected features based on defined threshold**

- rlength(weights*weights*ranked_weights_negative$rna)‘ RNAs have weights below -0.3 in factor 6
- 327 proteins have weights below -0.3 in factor 6

Some driving features may overlap between transcriptomics and proteomics

```
intersect(weights$weights$ranked_weights_negative$rna,weights$weights$ranked_weights_negative$protein)
```

```
##  [1] "ANXA1"   "B2M"     "ICAM5"   "COL18A1" "S100A10" "C4A"     "GFAP"
##  [8] "RLBP1"   "ITPR1"   "S100A4"  "GJA1"    "VIM"     "PRELP"   "CLIC1"
## [15] "AKAP5"   "FLNA"    "HSPG2"   "CAPS"    "AHNAK"   "COL6A2"  "CAPG"
## [22] "TAGLN"   "FABP7"   "C1QA"    "MYL9"    "F13A1"   "C3"      "COL14A1"
## [29] "OGN"     "ANXA2"   "C1QC"    "CRABP1"  "DCN"     "COL1A1"  "COL6A3"
## [36] "CALB2"
```

- 13 features overlap between omic layers

#### Multi-omics network

- Here we choose a moderate correlation threshold to draw edges between omic features (0.4)
- The same analysis can be repeated with an increased correlation threshold, which will yield more modules of strongly co-expressed features.

```
Weights_down_n=multiomics_network(multiassay=multiomics_object,
                           list=weights$weights$ranked_weights_negative,
                           correlation_threshold =0.4,
                           filter_string_50= TRUE)
```

##### Community detection within multi-omic network

The next step of the analysis is to find densily co-expressed communities of proteins and RNAs within the multi-omics network.

This function tries to find densely connected subgraphs in a graph by calculating the leading non-negative eigenvector of the modularity matrix of the graph.

```
communities <- communities_network(igraph=Weights_down_n$graph,
                                    community_detection='leading_eigen')
```

```
df <- purrr::map_dfr(
    .x = communities$communities,
    .f = ~ tibble::enframe(
      x = .x,
      name = NULL,
      value = "feature"
    ),
    .id = "community"
  )
```

- 5 communities are detected in the multi-omics network.
- Most communities are omic dependent, since generally there is more intra omic correlation than inter omic correlation

We can look at each communities:

##### Community 1

```
community_1=community_graph(igraph=Weights_down_n$graph,
                            community_object=communities$community_object,
                            community=1)
interactive_network(igraph=community_1$graph,communities=FALSE)
```

##### Community 4

```
community_4=community_graph(igraph=Weights_down_n$graph,
                            community_object=communities$community_object,
                            community=4)
interactive_network(igraph=community_4$graph,communities=FALSE)
```

##### Community hubs

- Features are more or less connected in the network, represented by their hub score:

```
print(sort(community_4$hubs, decreasing = TRUE))
```

```
 SLC1A5_rna    TBXAS1_rna  ARHGAP30_rna     SASH3_rna     LAIR1_rna
  1.00000000    0.99373217    0.98642752    0.98435469    0.98235011
  NCKAP1L_rna     ITGB2_rna       SLA_rna     RBM47_rna       SYK_rna
  0.98067378    0.97961400    0.97686353    0.97651079    0.96915836
    ALOX5_rna     HCLS1_rna    TYROBP_rna     IKZF1_rna     CD163_rna
  0.96356390    0.96296850    0.95458866    0.95094152    0.94339102
    C3AR1_rna        C3_rna      CD74_rna     CSF1R_rna   HLA-DRA_rna
  0.94277432    0.93651927    0.93628180    0.93554612    0.93182955
  SIGLEC9_rna    IL10RA_rna    MS4A4A_rna    ADORA3_rna      FGD2_rna
  0.92328733    0.92292107    0.91909216    0.91404692    0.91109604
```

```
  LAPTM5_rna     STAB1_rna       HCK_rna     PTAFR_rna      CD84_rna
  0.90461333    0.90405869    0.90396388    0.90011898    0.89988775
   CYTH4_rna     MS4A7_rna    SIRPB2_rna     RUNX1_rna    SH3TC1_rna
  0.89827538    0.89250311    0.89203385    0.89173034    0.89082576
    CD37_rna      IL18_rna    STEAP3_rna       FYB_rna      MSR1_rna
  0.88578012    0.88558732    0.88503274    0.87957245    0.87914404
    C1QC_rna     DOCK8_rna    CMKLR1_rna     VSIG4_rna      TLR5_rna
  0.87834137    0.87516337    0.87434232    0.87052433    0.86669267
   TREM2_rna      SCIN_rna      BTG2_rna   SIGLEC8_rna      C1QA_rna
  0.86641949    0.86251896    0.85755337    0.85515538    0.84538301
   TGFBI_rna    PIK3R5_rna   ADAMTS1_rna    CLEC7A_rna     S1PR3_rna
  0.84536757    0.84233627    0.83583036    0.83423436    0.82535731
 TMEM119_rna      CTSS_rna     CLIC1_rna      TLR7_rna    IQGAP2_rna
  0.82196903    0.82151641    0.82109966    0.81735518    0.80021262
    FGL2_rna     KLHL6_rna     LTBP2_rna  SIGLEC10_rna   HLA-DMB_rna
  0.80014392    0.79227627    0.78811550    0.78656997    0.78629965
  COL8A1_rna      UCP2_rna     CIITA_rna      C1QB_rna    FBLIM1_rna
  0.78532943    0.78043171    0.77845326    0.77178803    0.76323399
HLA-DQA1_rna    SLC2A5_rna   ALOX15B_rna   SLC37A2_rna     FKBP5_rna
  0.75994557    0.75408226    0.75017993    0.74853906    0.74461447
    THBD_rna  HLA-DPA1_rna    SELPLG_rna    RNASE6_rna    MS4A6A_rna
  0.73764817    0.73642525    0.73573903    0.72218947    0.70762774
    CCR1_rna   FAM129A_rna      CD14_rna        C7_rna      IER3_rna
  0.70440255    0.70378491    0.70233392    0.70213043    0.68454576
    CD53_rna    S100A4_rna    PDLIM1_rna    GPRC5A_rna     PARVG_rna
  0.67813768    0.67782265    0.67106666    0.66807648    0.66113899
    SGK1_rna     GPR34_rna     ITGAX_rna     SPHK1_rna  C11orf96_rna
  0.65618938    0.65134262    0.65084388    0.64913399    0.64352477
   ADAM28_rna    PCDH18_rna    CYP1B1_rna      NGFR_rna     THBS1_rna
  0.63338782    0.62817867    0.62545763    0.62241502    0.61730883
   F13A1_rna   SIGLEC1_rna    IGFBP4_rna        F5_rna    MS4A14_rna
  0.59992031    0.56682474    0.56215001    0.55344578    0.55266215
    MRC1_rna     MEDAG_rna VAT1L_protein     SH2D6_rna      APOD_rna
  0.54694702    0.52958781    0.51949931    0.51491910    0.45485914
   HMOX1_rna     CXCL2_rna PTBP2_protein    AKR1C2_rna ACOT1_protein
  0.45093102    0.42784352    0.38379233    0.36378187    0.32706950
    MMP9_rna      LIPG_rna     ACKR1_rna
  0.29932805    0.10494025    0.07805484
```

- Top gene in module 4 is SLC1A5_rna
- TREM2 or CLU, known AD risk genes expressed on microglia are present here
- From prior knowledge, many of the genes in module 4 are related to microglial activation

#### Cell type enrichment of modules

Here we proceed with a `EWCE` analysis to check if modules are enriched in specific cell types.

```
cell_type <- cell_type_enrichment(
    multiassay =multiomics_object,
    communities = lapply(communities$communities,function(x){sub("\\_.*", "",x )}),
    ctd= ctd
  )

cell_type$plots
```

```
## $`1`
```

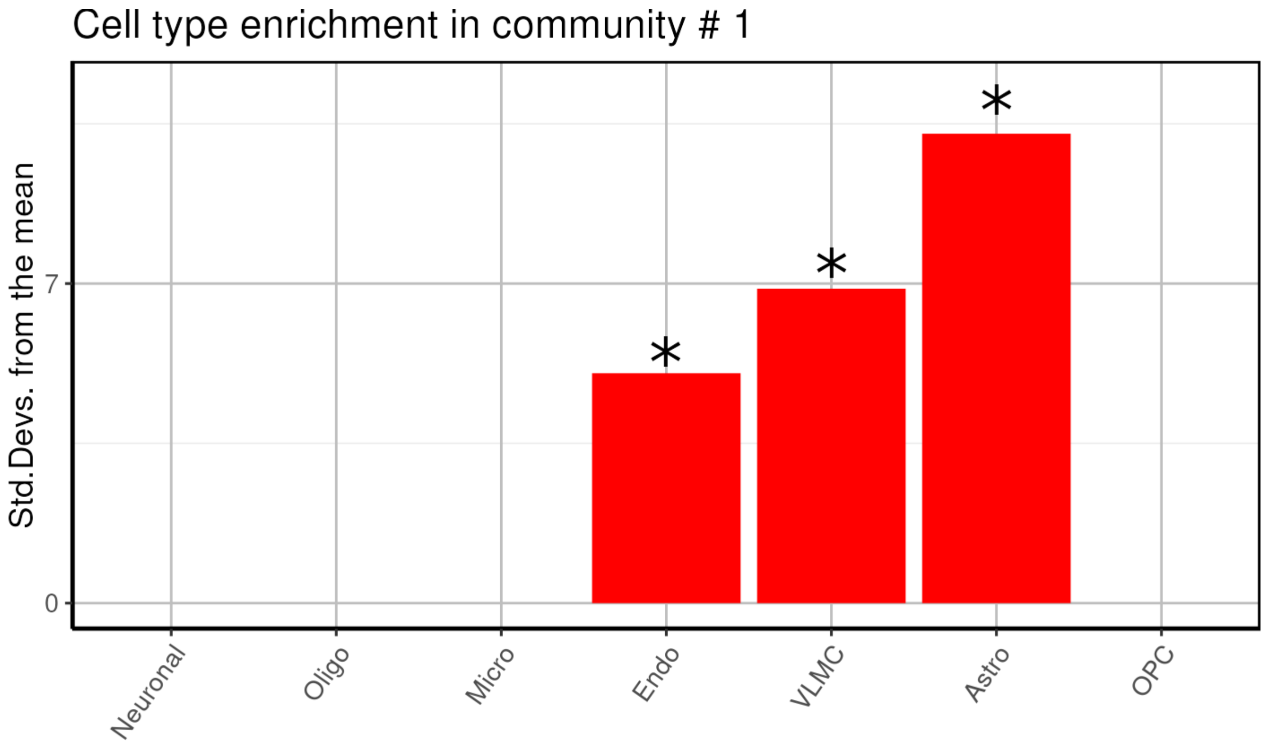
Cell type enrichment in community # 1
Std.Devs. from the mean
7
0
*
*
*
Neuronal
Oligo
Micro
Endo
VLMC
Astro
OPC

```
##
## $`2`
```

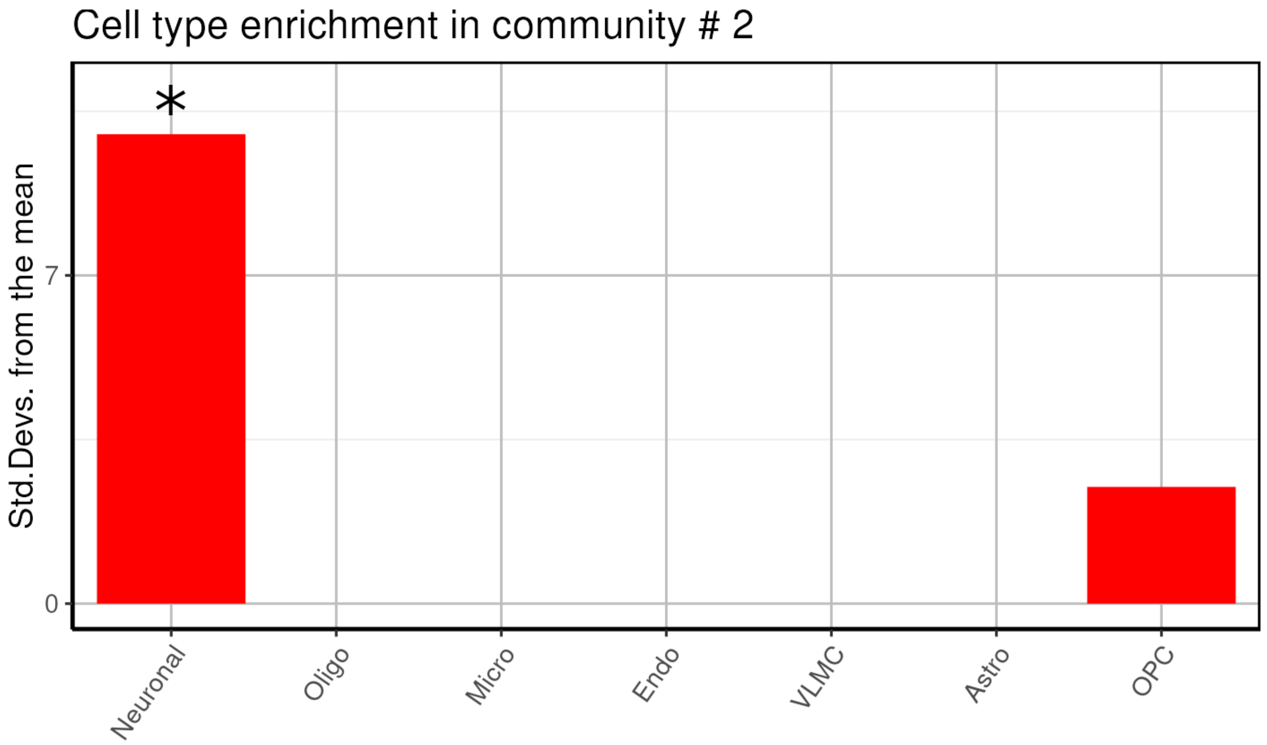
Cell type enrichment in community # 2
Std.Devs. from the mean
7
0
*
Neuronal
Oligo
Micro
Endo
VLMC
Astro
OPC

```
##
## $`3`
```

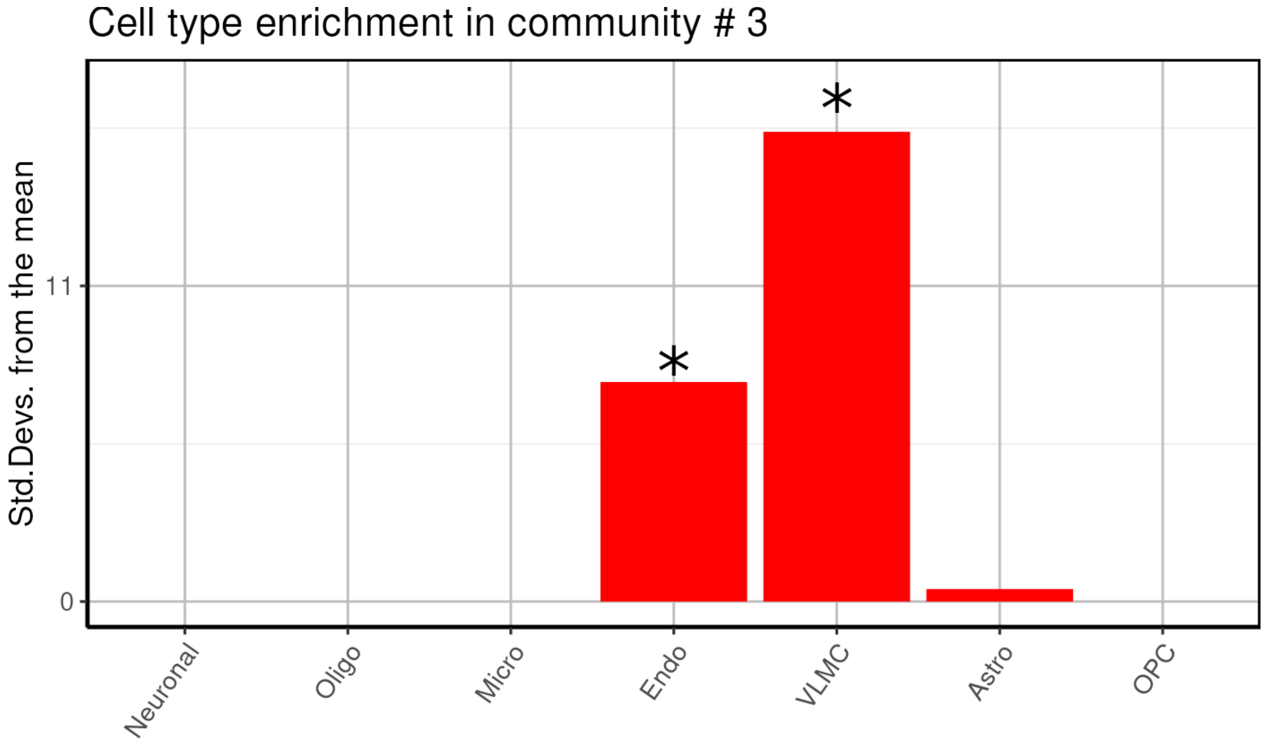
Cell type enrichment in community # 3
Std.Devs. from the mean
11
0
*
*
Neuronal
Oligo
Micro
Endo
VLMC
Astro
OPC

```
##
## $`4`
```

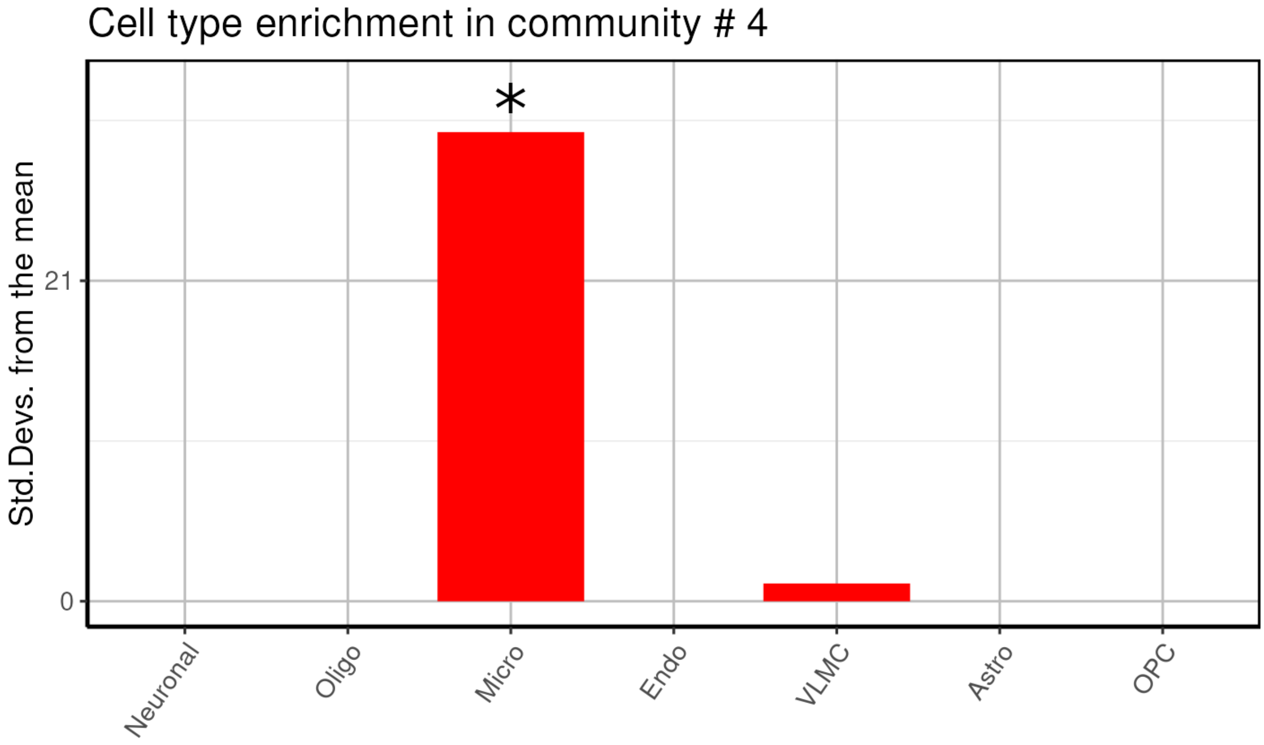
Cell type enrichment in community # 4
Std.Devs. from the mean
21
0
*
Neuronal
Oligo
Micro
Endo
VLMC
Astro
OPC

```
##
## $`5`
```

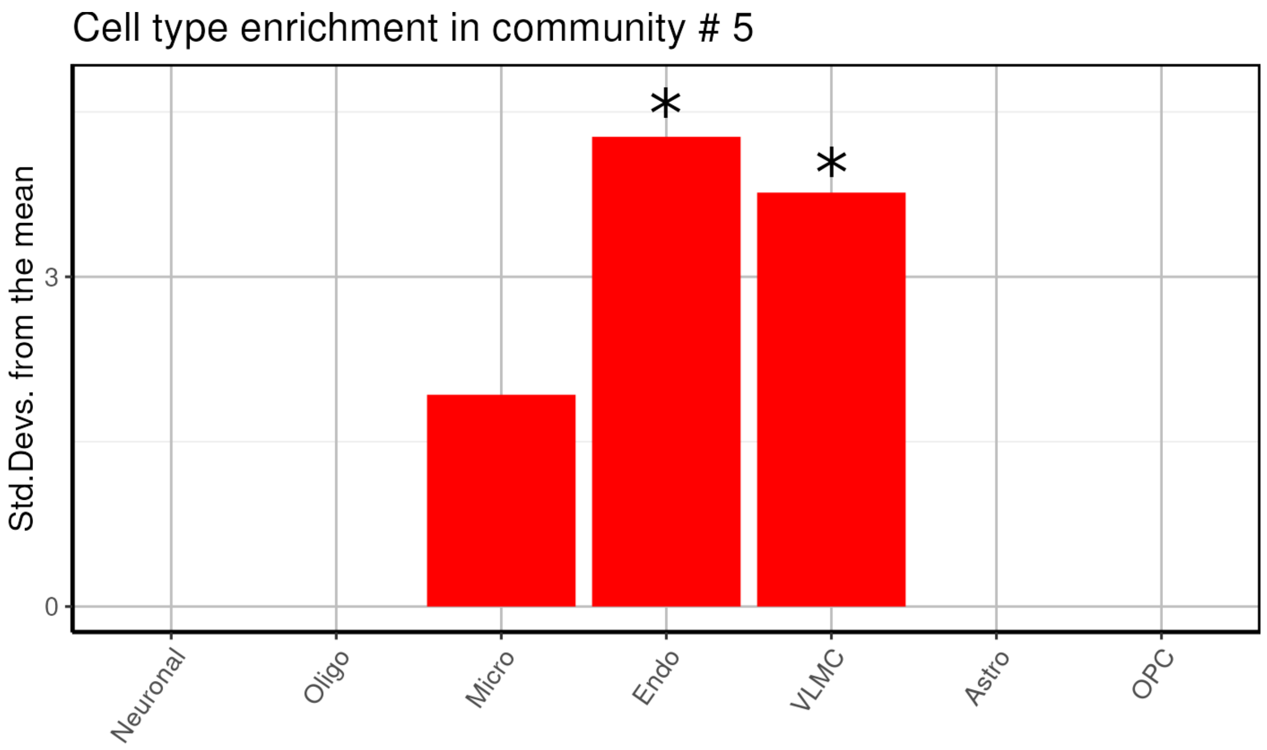

- We find one neuronal, one microglial and two undefined glial enriched modules.

##### Pathway enrichment analysis

```
communities_l = lapply(communities$communities,function(x){sub("\\_.*", "",x )})

enrichment1=pathway_analysis_enrichr(communities_l$`1`,plot=5)
```

```
## Welcome to enrichR
## Checking connection ...

## Enrichr ... Connection is Live!
## FlyEnrichr ... Connection is available!
## WormEnrichr ... Connection is available!
## YeastEnrichr ... Connection is available!
## FishEnrichr ... Connection is available!

## Uploading data to Enrichr... Done.
##   Querying GO_Molecular_Function_2021... Done.
##   Querying GO_Cellular_Component_2021... Done.
##   Querying GO_Biological_Process_2021... Done.
##   Querying WikiPathways_2021_Human... Done.
##   Querying Reactome_2016... Done.
##   Querying KEGG_2021_Human... Done.
##   Querying MSigDB_Hallmark_2020... Done.
##   Querying BioCarta_2016... Done.
## Parsing results... Done.
```

```
## i Output is returned as a list!
```

```r
enrichment2=pathway_analysis_enrichr(communities_l$`2`,plot=5)
```

```
## Welcome to enrichR
## Checking connection ...
## Enrichr ... Connection is Live!
## FlyEnrichr ... Connection is available!
## WormEnrichr ... Connection is available!
## YeastEnrichr ... Connection is available!
## FishEnrichr ... Connection is available!

## Uploading data to Enrichr... Done.
##   Querying GO_Molecular_Function_2021... Done.
##   Querying GO_Cellular_Component_2021... Done.
##   Querying GO_Biological_Process_2021... Done.
##   Querying WikiPathways_2021_Human... Done.
##   Querying Reactome_2016... Done.
##   Querying KEGG_2021_Human... Done.
##   Querying MSigDB_Hallmark_2020... Done.
##   Querying BioCarta_2016... Done.
## Parsing results... Done.

## i Output is returned as a list!
```

```r
enrichment3=pathway_analysis_enrichr(communities_l$`3`,plot=5)
```

```
## Welcome to enrichR
## Checking connection ...
## Enrichr ... Connection is Live!
## FlyEnrichr ... Connection is available!
## WormEnrichr ... Connection is available!
## YeastEnrichr ... Connection is available!
## FishEnrichr ... Connection is available!

## Uploading data to Enrichr... Done.
##   Querying GO_Molecular_Function_2021... Done.
##   Querying GO_Cellular_Component_2021... Done.
##   Querying GO_Biological_Process_2021... Done.
##   Querying WikiPathways_2021_Human... Done.
##   Querying Reactome_2016... Done.
##   Querying KEGG_2021_Human... Done.
##   Querying MSigDB_Hallmark_2020... Done.
##   Querying BioCarta_2016... Done.
## Parsing results... Done.

## i Output is returned as a list!
```

```r
enrichment4=pathway_analysis_enrichr(communities_l$`4`,plot=5)
```

```
## Welcome to enrichR
## Checking connection ...
## Enrichr ... Connection is Live!
## FlyEnrichr ... Connection is available!
## WormEnrichr ... Connection is available!
## YeastEnrichr ... Connection is available!
## FishEnrichr ... Connection is available!

## Uploading data to Enrichr... Done.
```

```
##   Querying GO_Molecular_Function_2021... Done.
##   Querying GO_Cellular_Component_2021... Done.
##   Querying GO_Biological_Process_2021... Done.
##   Querying WikiPathways_2021_Human... Done.
##   Querying Reactome_2016... Done.
##   Querying KEGG_2021_Human... Done.
##   Querying MSigDB_Hallmark_2020... Done.
##   Querying BioCarta_2016... Done.
## Parsing results... Done.

## i Output is returned as a list!
```

```
enrichment5=pathway_analysis_enrichr(communities_l$`5`,plot=5)
```

```
## Welcome to enrichR
## Checking connection ...
## Enrichr ... Connection is Live!
## FlyEnrichr ... Connection is available!
## WormEnrichr ... Connection is available!
## YeastEnrichr ... Connection is available!
## FishEnrichr ... Connection is available!

## Uploading data to Enrichr... Done.
##   Querying GO_Molecular_Function_2021... Done.
##   Querying GO_Cellular_Component_2021... Done.
##   Querying GO_Biological_Process_2021... Done.
##   Querying WikiPathways_2021_Human... Done.
##   Querying Reactome_2016... Done.
##   Querying KEGG_2021_Human... Done.
##   Querying MSigDB_Hallmark_2020... Done.
##   Querying BioCarta_2016... Done.
## Parsing results... Done.

## i Output is returned as a list!
```

```
enrichment1$plot$Reactome_2016+theme(axis.text=element_text(size=12),
                                     axis.title=element_text(size=12),
                                     strip.text = element_text(size = 0, margin = margin()),
                                     legend.title = element_text(size=9))
```

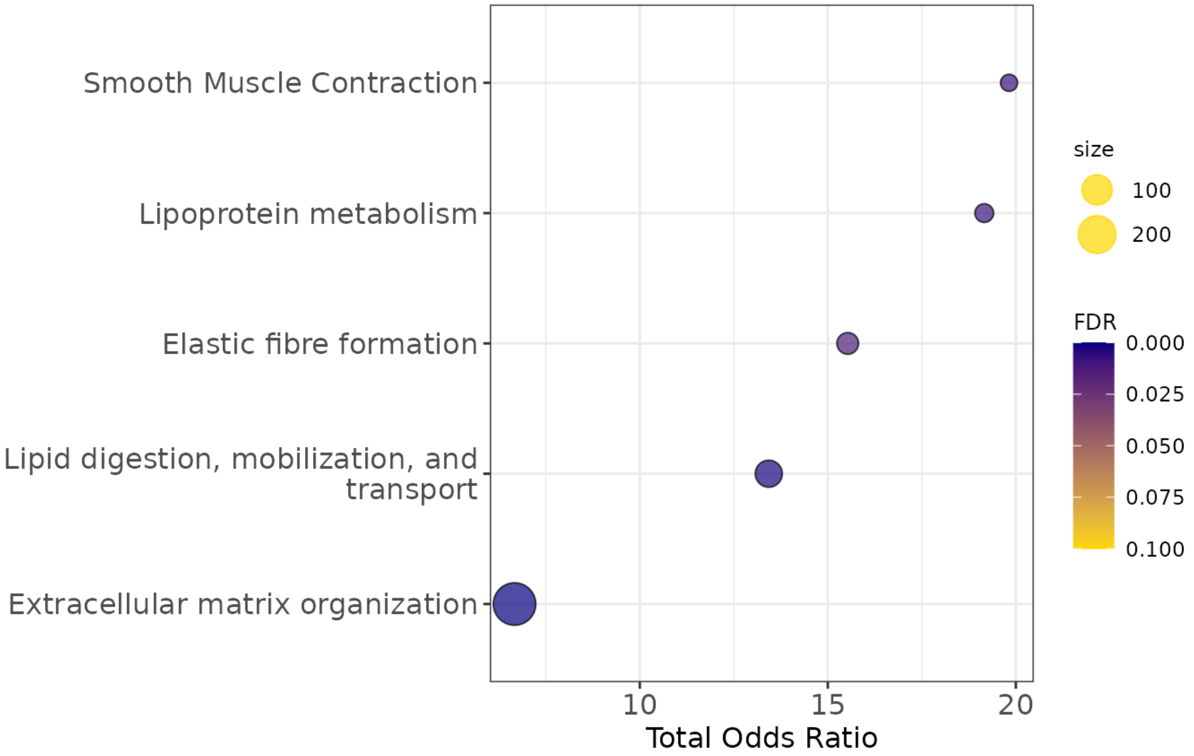

```
enrichment2$plot$Reactome_2016+theme(axis.text=element_text(size=12),
                                     axis.title=element_text(size=12),
                                     strip.text = element_text(size = 0, margin = margin()),
                                     legend.title = element_text(size=9))
```

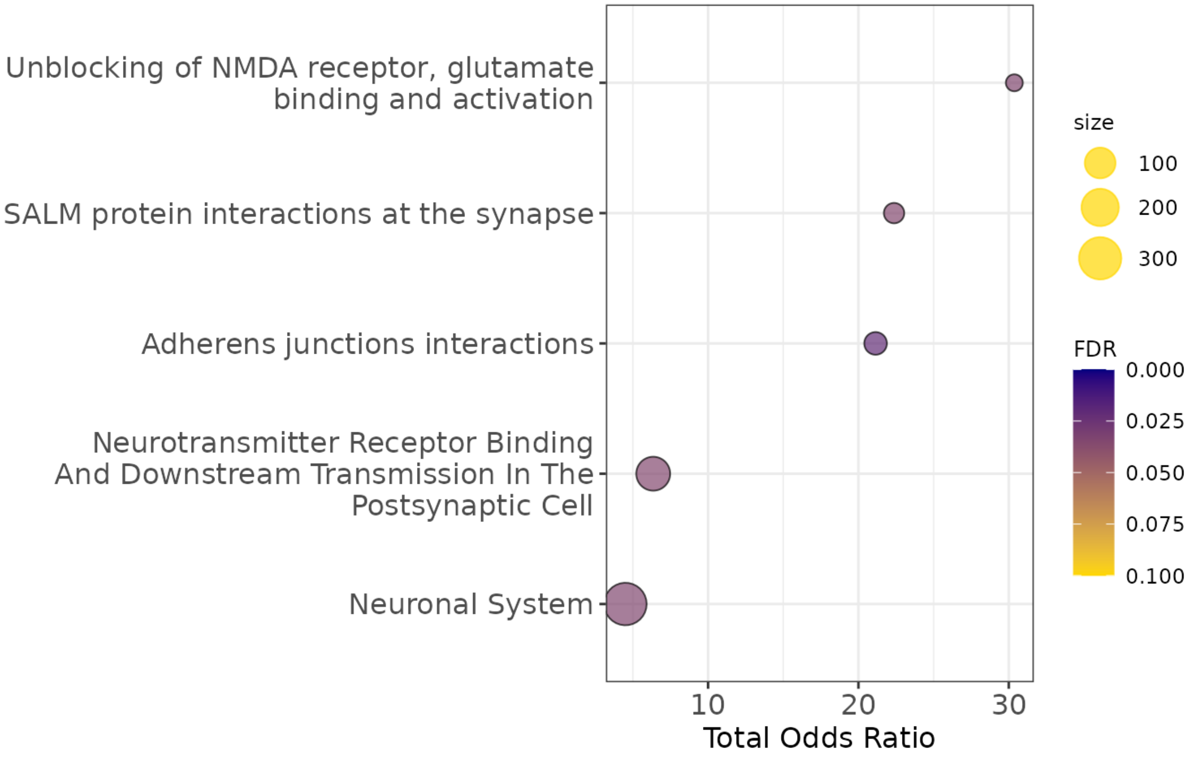

```
enrichment3$plot$Reactome_2016+theme(axis.text=element_text(size=12),
                                     axis.title=element_text(size=12),
                                     strip.text = element_text(size = 0, margin = margin()),
                                     legend.title = element_text(size=9))
```

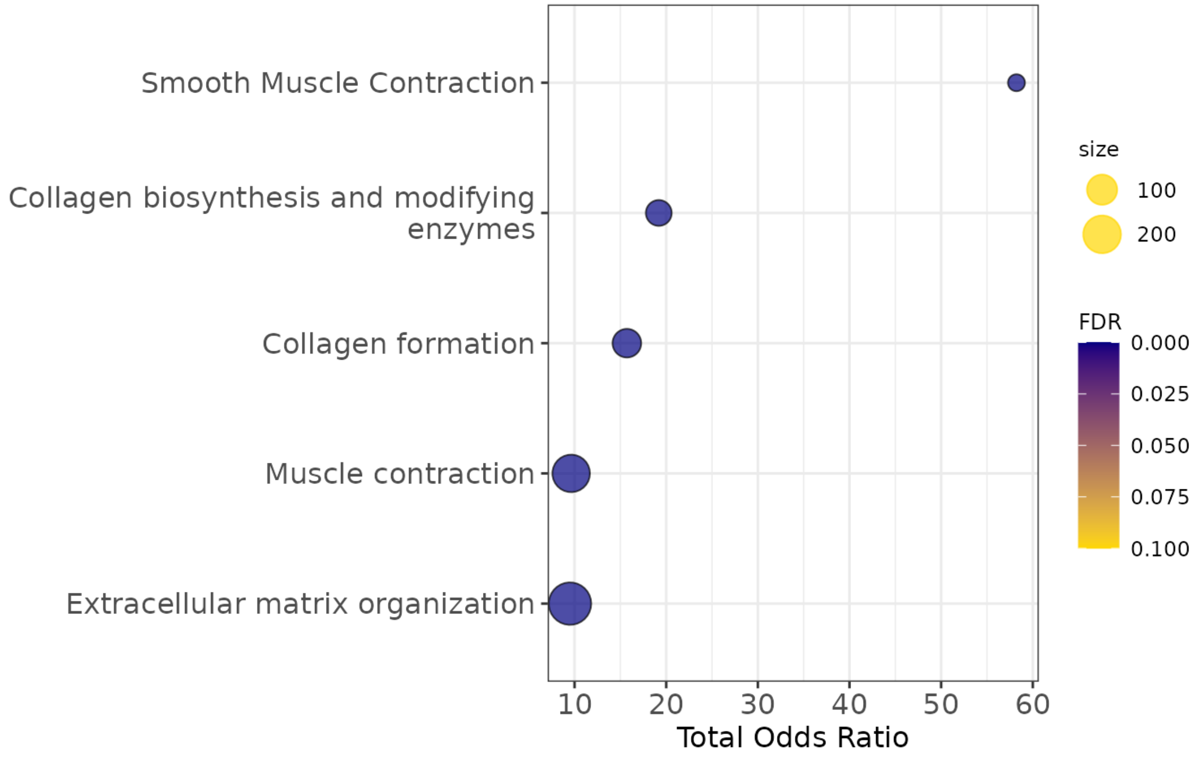

```
enrichment4$plot$Reactome_2016+theme(axis.text=element_text(size=12),
                                     axis.title=element_text(size=12),
                                     strip.text = element_text(size = 0, margin = margin()),
                                     legend.title = element_text(size=9))
```

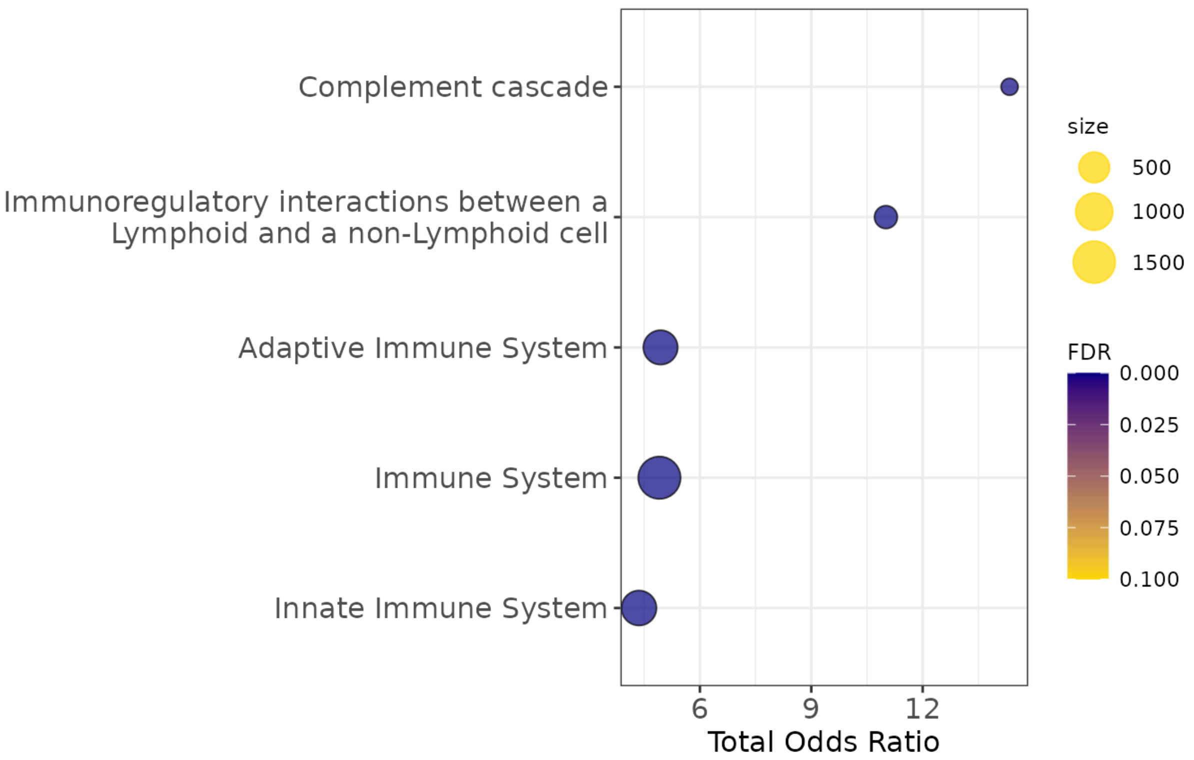

```
enrichment5$plot$Reactome_2016+theme(axis.text=element_text(size=12),
                                     axis.title=element_text(size=12),
                                     strip.text = element_text(size = 0, margin = margin()),
                                     legend.title = element_text(size=9))
```

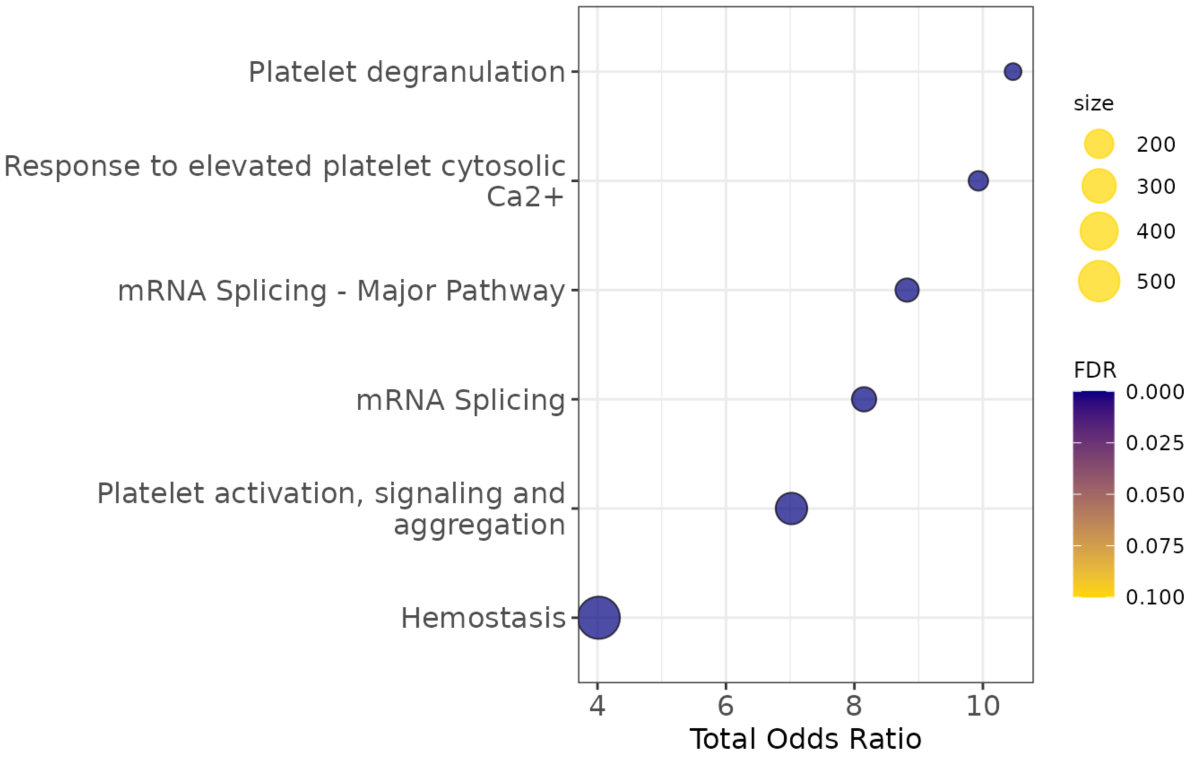

##### Modules correlation to neuropathology

`multiomics_modules` compute the module eigen value (PC1 of scaled transcriptomics/proteomics expression) and correlates it to chosen covariates

```
modules=multiomics_modules(multiassay=multiomics_object,
                           metadata=MOFA2::samples_metadata(integrated_object),
                           covariates=c('inferred_pseudotime','PHF1','amyloid','pTau'),
                           communities=communities$communities,
                           filter_string_50=TRUE)
```

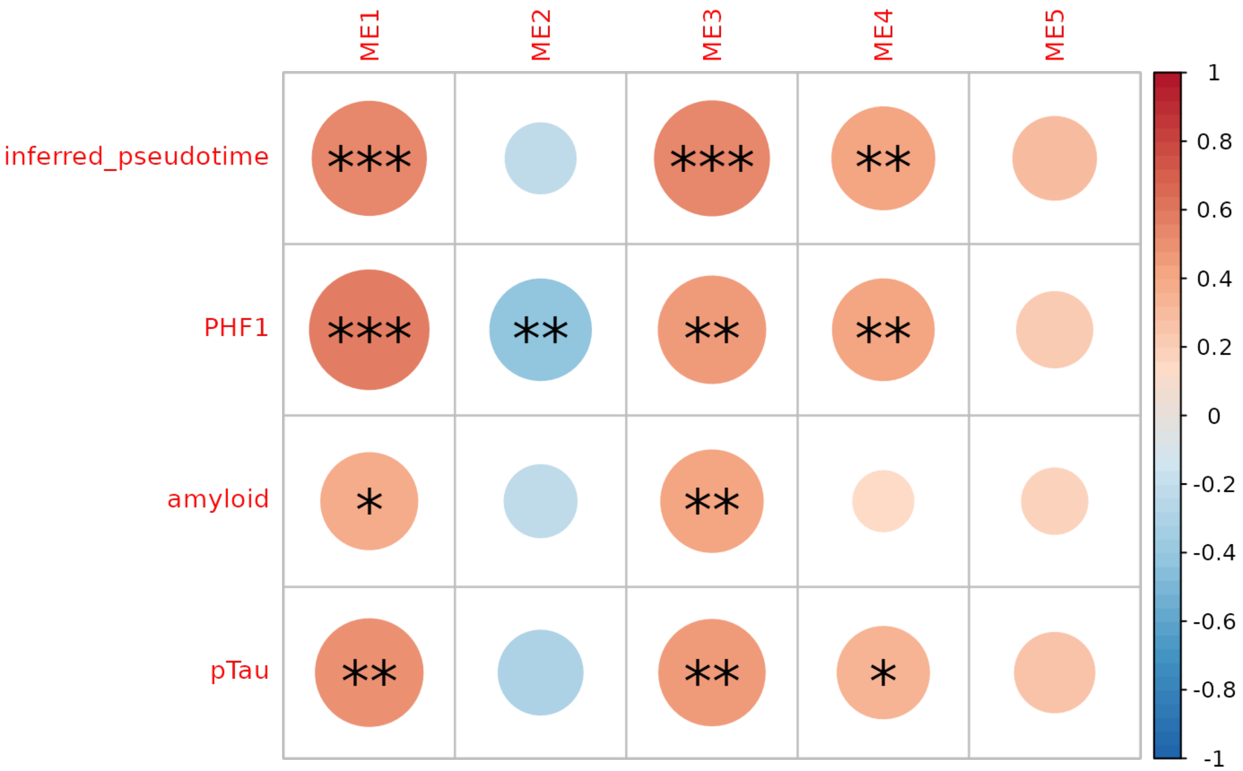

##### Transcription Factor - target gene enrichment

```
TF=Transcription_Factor_enrichment(communities=communities$communities,
                                   weights=weights,
                                   TF_gmt=TF_fp,
                                   threshold = 0.2,
                                   direction = "negative")

GeneOverlap::drawHeatmap(TF,what = c("odds.ratio"), log.scale = T, adj.p = F, cutoff = .05,
                         ncolused = 7, grid.col = c("Blues"), note.col = "red")
```

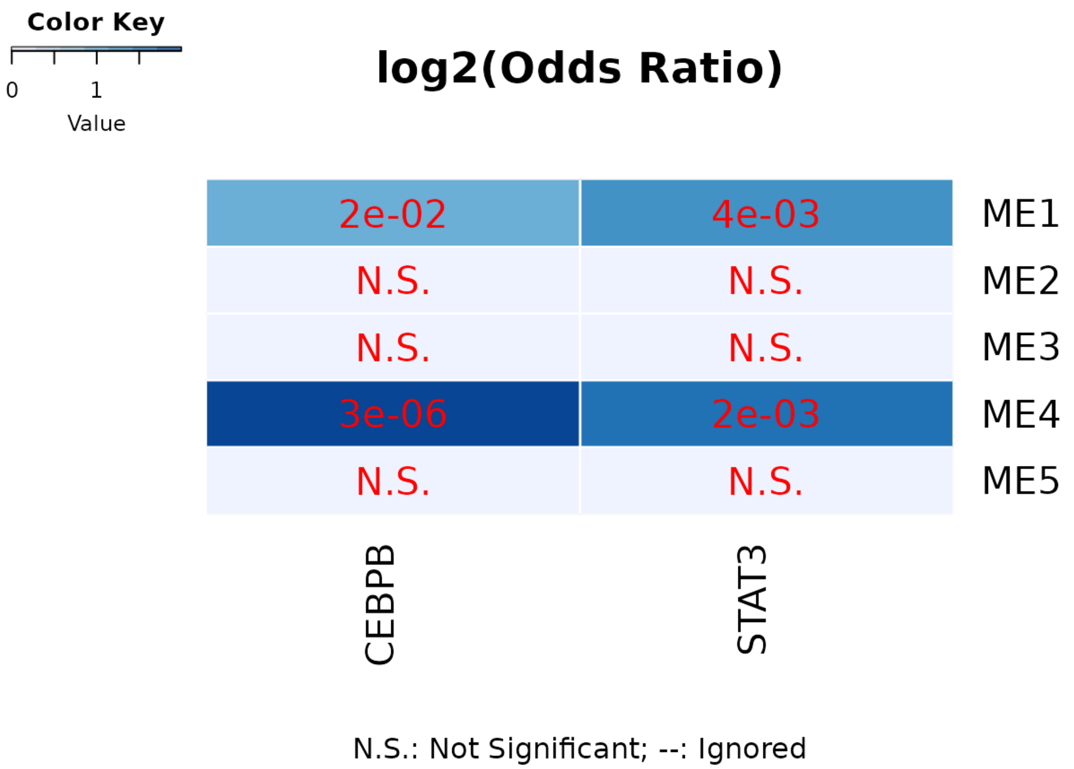

##### TF-target gene circos plot

```
circos_TF(TF)
```

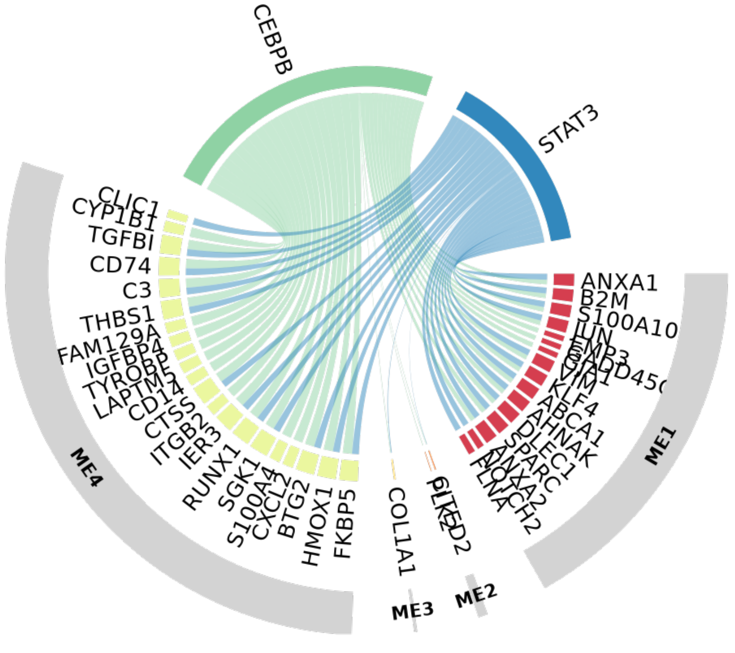

##### Pseudotime ordering of multi-omics modules

- Here we related each module eigen value to the pseudotime inferred from MOFA integration emmbeddings (Factors 6/8)

```
plot_module_trajectory(modules=modules,
                       covariates=c('amyloid','pTau','PHF1'),
                       plot_modules=c("ME1","ME2","ME3","ME4"))
```

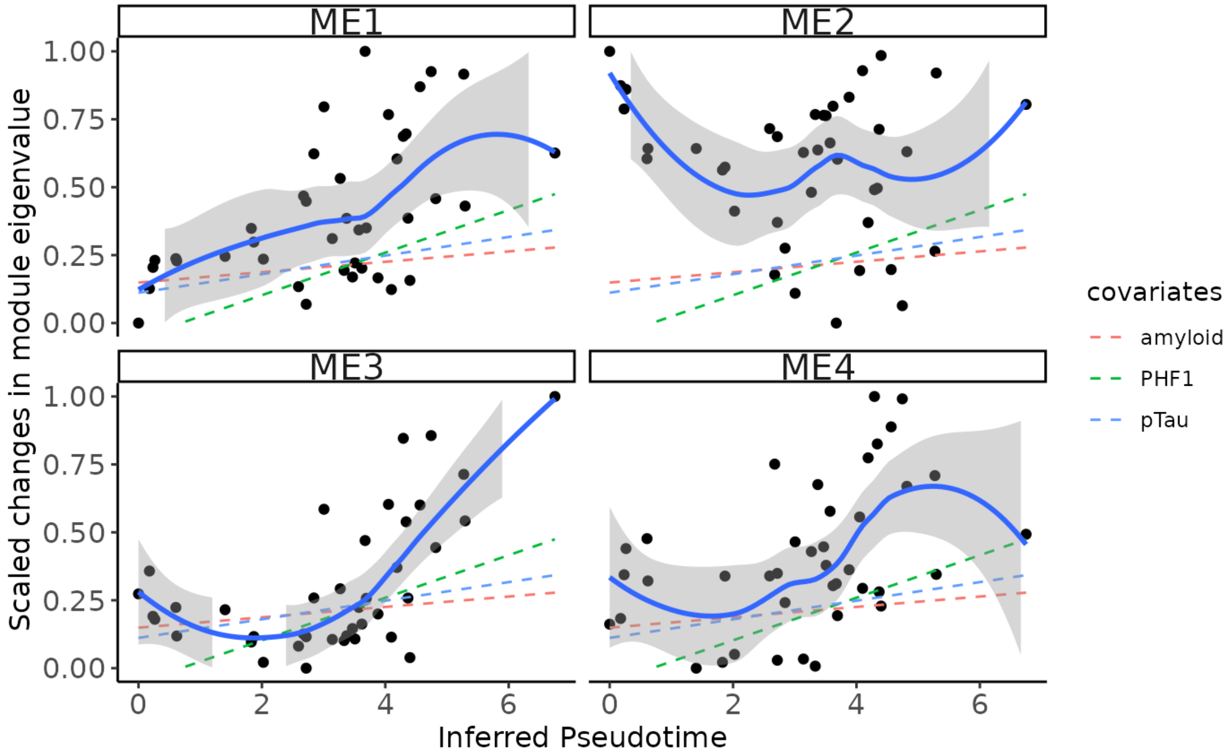
